# Cortical projection neurons with distinct axonal connectivity employ ribosomal complexes with distinct protein compositions

**DOI:** 10.1101/2024.12.22.629968

**Authors:** Tien Phuoc Tran, Bogdan Budnik, John E. Froberg, Jeffrey D. Macklis

## Abstract

Diverse subtypes of cortical projection neurons (PN) form long-range axonal projections that are responsible for distinct sensory, motor, cognitive, and behavioral functions. Translational control has been identified at multiple stages of PN development, but how translational regulation contributes to formation of distinct, subtype-specific long-range circuits is poorly understood. Ribosomal complexes (RCs) exhibit variations of their component proteins, with an increasing set of examples that confer specialized translational control. Here, we directly compare the protein compositions of RCs *in vivo* from two closely related cortical neuron subtypes–cortical output “subcerebral PN” (SCPN) and interhemispheric “callosal PN” (CPN)– during establishment of their distinct axonal connectivity. Using retrograde labeling of subtype-specific somata, purification by fluorescence-activated cell sorting, ribosome immunoprecipitation, and ultra-low-input mass spectrometry, we identify distinct protein compositions of RCs from these two subtypes. Strikingly, we identify 16 associated proteins reliably and exclusively detected only in RCs of SCPN. 11 of these proteins have known interaction with components of ribosomes; we further validated ribosome interaction with protein kinase C epsilon (PRKCE), a candidate with roles in synaptogenesis. PRKCE and a subset of SCPN-specific candidate ribosome-associated proteins also exhibit enriched gene expression by SCPN. Together, these results indicate that ribosomal complexes exheq]ibit subtype-specific protein composition in distinct subtypes of cortical projection neurons during development, and identify potential candidates for further investigation of function in translational regulation involved in subtype-specific circuit formation.

## Introduction

The brain’s ability to perform distinct sensory, motor, cognitive, and behavioral functions relies critically on the proper development of diverse neuronal subpopulations, and establishment of their distinct connectivity. In the mammalian cerebral cortex, excitatory glutamatergic projection neurons (PN) originate from dorsal pallial progenitors and migrate radially to form the cortex in an inside-out manner, with early-born neurons occupying deeper layers and later-born neurons settling in more superficial layers^1^. Distinct subtypes of PN then extend long-range axonal projections to specific targets, forming circuits that control motor function, sensory perception, cognition, and associative behavior.

We focus on two exemplar PN subtypes, defined by their primary axonal connectivity. Subcerebral PN (SCPN), which include corticospinal neurons, are a distinct subtype of cortical output neurons that comprises the entire population of neurons that project from the cortex to the brainstem and spinal cord^1,2^. SCPN reside predominantly in layer V, and control voluntary and dextrous motor function. SCPN of motor and non-motor types are specifically vulnerable to degeneration in amyotrophic lateral sclerosis (ALS)^3,4^ and related frontotemporal dementia (FTD)^5,6^, respectively. In contrast, callosal PN (CPN), with ∼80% in cortical layer II/III and ∼20% in layer V, extend axons through the corpus callosum (CC) and connect generally homotopic areas between hemispheres^7^. CPN are essential for high-level associative and integrative functions, and are involved in autism spectrum disorders, intellectual disability, and other neuropsychiatric disorders.

SCPN and CPN are developmentally closely related cortical PN, and must both establish shared properties of long-projection, glutamatergic PN while building quite distinct axonal connectivity during development, so relatively subtle molecular distinctions between these subtypes likely underlie their subtype-specific circuit formation. In the past two decades, transcriptional analyses of SCPN, CPN, and a few other well-studied subtypes have identified key transcriptional regulatory controls over axonal connectivity (reviewed in^1^). Translational regulatory control during cortical development is less well investigated, but is likely an important additional layer of dynamic and spatial regulatory control.

Translational regulation controls when, where, and how proteins are made, and multiple lines of evidence support such functions in cortical development. In early corticogenesis, progenitors and newly born neurons often co-express multiple transcription factors specifying alternative subtype identities at the mRNA level, but repress their translation, likely to ensure temporal precision of TF expression^8,9^. As neurogenesis progresses from production of early-born neurons to production of later-born neurons, there are shifts in subsets of transcripts undergoing translation, regulated by RNA-binding proteins including ELAVL4^10^, ELAVL1^11^, and CELF1^12^ as well as by ribosomal proteins RPL7/uL30 and RPL10/uL16^13^. Later-born neurons exhibit higher global translational rates than their early-born counterparts^14^.

During later stages, as PN extend long-range axonal projections, specific transcripts are trafficked to axonal growth cones^15,16^, and *in vitro* work has revealed that local translation functions importantly in axonal growth and guidance^17–20^.

Despite these insights that underscore the importance of translational control, understanding is limited as to how this regulation is specialized in distinct projection neuron sub-types, especially while subtype-specific long-range circuitry is being established. In addition to selection of mRNAs, little is known about subsequent steps of translation, including mechanisms governing subcellular-organellar localization of protein production, elongation rate, as well as co-translational folding and modifications. Moreover, few molecular regulators of translational control, and protein synthesis in general, have been identified in cerebral cortex *in vivo*, let alone in specific subtypes.

To investigate subtype-specificity of translational control *in vivo*, our lab developed two complementary approaches. In previously reported work, Froberg *et al*. developed and applied “nanoRibo-Seq”, an ultra-low-input ribosome profiling approach, to enable subtype-specific investigation of transcriptome-wide translational efficiency from purified somata^21^. While this work at one level confirmed the expected high correlation between broad SCPN and CPN translational patterns, it notably identified about four dozen mRNAs with significantly differential translational efficiencies between SCPN and CPN^21^. Strikingly, the analyses also quite unexpectedly identified extensive translation from upstream open reading frames (uORFs) for synapse-related genes for both subtypes^21^. This work reinforced the importance of subtype-specific translational regulation, strongly motivating investigation of subtype-specific molecular machinery that might underlie such differences.

Here, we directly investigate potential specialization of translational machinery: ribosomes and their associated complexes, collectively referred to here as “ribosomal complexes” (RCs).

Emerging evidence highlights the vast potential for specialization of ribosomes by modifying their protein composition of both core ribosomal proteins (RPs) and associated proteins^22^. Several RPs including RPL10A/uL1^23^, RPL38/eL38^23,24^, and RPS25/eS25^23^ are present only in subsets of ribosomes, and are found to preferentially bind subsets of transcripts. Ribosomes also serve as a hub for translation factors, RNA-binding and processing proteins^22,25^ and components of co-translational modification, folding, and complex assembly machinery^26^. Associated proteins play critical roles in translation, and are increasingly found to confer to ribosomes remarkably varied functional specialization^22,25^. Examples in neurobiology include: fragile X mental retardation protein (FMRP), which inhibits translation elongation of its target transcripts^27^; and p180, which localizes ribosomes to axonal ER tubules and recruits mRNAs encoding axonal membrane proteins^28^. Intriguingly, the axon guidance receptor DCC inhibits translation via binding to ribosomes, which are released upon DCC binding to its ligand, Netrin^29^.

These examples indicate that specific protein compositions of ribosomal complexes result in distinct regulation of translation. Varying core RP and associated cytoplasmic protein composition is potentially a mechanism to rapidly tune translation, which might be especially advantageous for post-mitotic neurons as they undergo distinct dynamic stages of development while avoiding the energetic demands of producing new RCs. Thus, in the context of subtype-specific circuit development, we investigate here the hypothesis that distinct subtypes of PN might possess specialized RCs with unique protein composition during development, enabling precise translational control underlying specific circuit formation.

To test this hypothesis, we directly compared RCs isolated from purified somata of mouse SCPN and CPN at post-natal day 3 (P3) *in vivo*, during which both subtypes are establishing precise long-range axonal connectivity. To directly investigate the protein compositions of RCs in these subtypes, we used a combination of retrograde labeling of circuit-specific somata, subtype-specific neuronal purification by fluorescence-activated cell sorting, ribosomal RNA (rRNA) immunoprecipitation, and ultra-low-input mass spectrometry (IP-MS).

We identify that SCPN and CPN have distinct ribosomal complexes (RCs), with 16 non-core proteins that are exclusively present in SCPN RCs and that span varied functions (including chaperones, metabolic enzymes, kinases, and an RNA-binding protein). We also find enrichment of RPS30/eS30 in CPN RCs, suggesting potential heterogeneity of core ribosome protein composition. For a subset of proteins specific to RCs of SCPN, we obtain evidence for their physical interaction with ribosomal components. Selecting protein kinase C-epsilon (PRKCE) for further verification based on its known roles in synaptogenesis, we find, e.g., that PRKCE reciprocally pulls down ribosomes in P3 cortex. Further, PRKCE and multiple proteins with validated ribosome interactions also exhibit enriched gene expression and concordantly more translation in SCPN compared to CPN. Together, these results both indicate subtype-specificity of ribosomal complexes between distinct neuronal subtypes, and identify several potential translational regulators that might function in unique ways in subtype-specific circuit development.

## Results

### Affinity purification of ribosome-associated complexes from purified SCPN and CPN *in vivo*

To investigate the protein composition of endogenous RCs from specific cortical PN subtypes *in vivo*, we combined retrograde labeling of circuit-specific neuronal somata for their purification by fluorescence-activated cell sorting (FACS) with rRNA IP-MS (Figure 1A). Retrograde labeling reliably labels and distinguishes distinct projection neuron subtypes based on their distinct combinations of axonal connectivity and soma position, a long-standing and well-accepted method that is found to cause minimal damage and not to interfere with continued neuronal development, and that has led to the identification of cardinal regulatory genes and molecules of axonal connectivity^2,30–36^. Here, dual retrograde labeling of SCPN and CPN at P1 in the same mice quite distinctly labels the appropriate, non-overlapping subpopulations of cortical PN by P3 (Figure 1B). In particular, fluorophore-conjugated cholera toxin B (CTB) injection into the right side of the corpus callosum prominently labels CPN in layer II/III of the left hemisphere. CTB injection into the left side of the cerebral peduncle appropriately labels SCPN somata in layer V of the left hemisphere. This direct validation of CPN vs. SCPN distinction and labeling specificity enabled us to employ one or the other single retrograde labeling strategy to purify either CPN or SCPN from separate mice for proteomic investigation.

**Figure 1.**
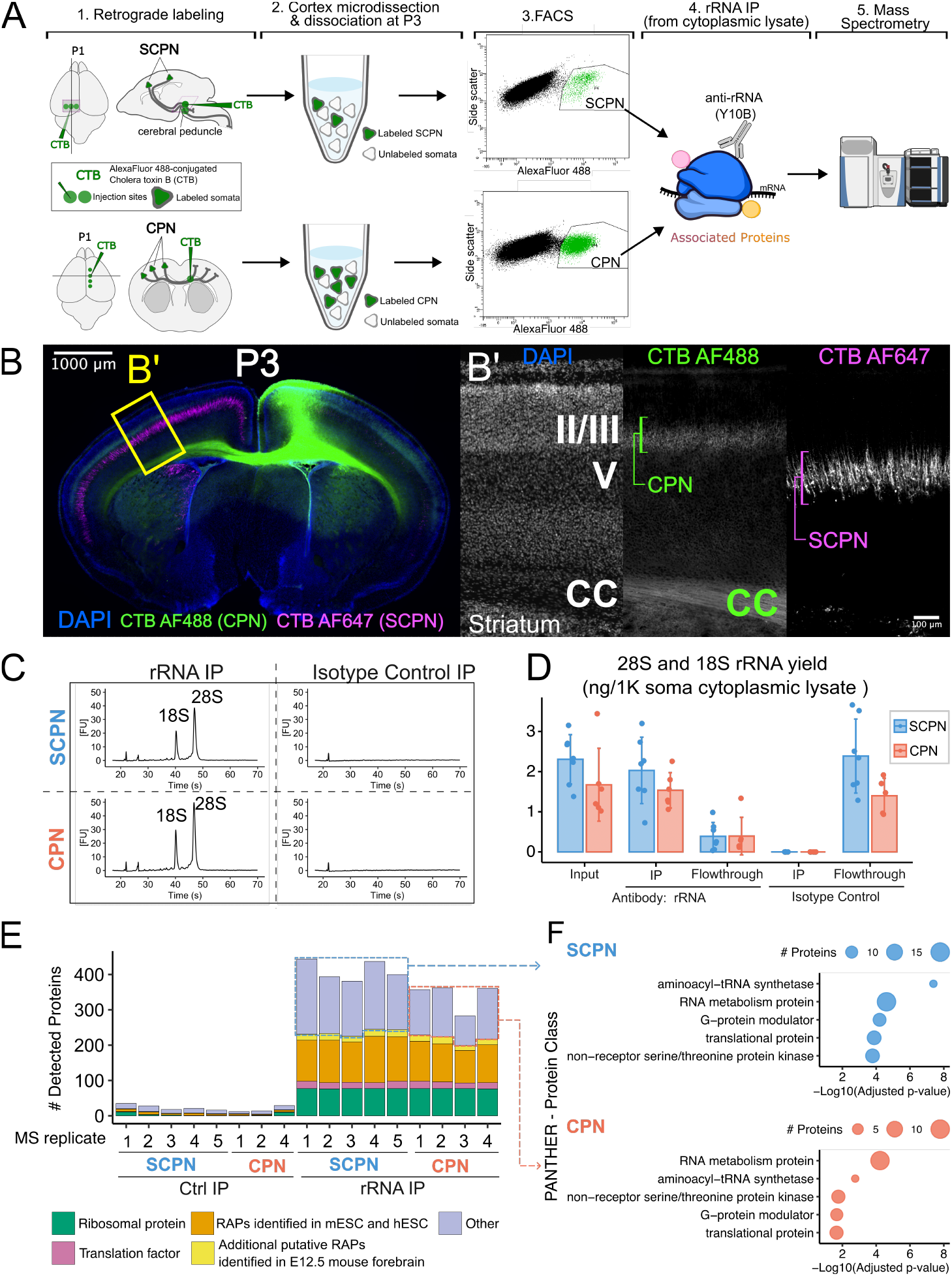
Affinity purification of ribosomal complexes from purified somata of SCPN and CPN *in vivo*. **(A)** Workflow to investigate ribosomal complexes from specific cortical projection neuron (PN) subtypes *in vivo*. **(B)** Dual retrograde labeling of CPN and SCPN at P1 in the same mice distinctly labels the appropriate, non-overlapping subpopulations of cortical PN by P3. Inset B’ shows the location of cortical layers II/III and V, striatum, and corpus callosum (CC). (n=4 biological replicates) **(C)** Bioanalyzer characterization shows specific recovery of rRNAs from both FACS-purified CPN and SCPN using rRNA IP. **(D)** Quantification of 28S and 18S rRNA recovery following IP with rRNA antibody or isotype control antibody in purified CPN and SCPN. Individual dots represent biological replicates, with bars showing mean values and error bars indicating standard deviation. **(E)** Mass spectrometry analysis shows enrichment of most ribosomal proteins (76-77 out of ∼80 known proteins), translation factors, and known ribosome-associated proteins (RAPs) in rRNA IP samples compared to control IP samples from both SCPN and CPN. RAPs were reported in Simsek *et al*. Cell 2017^25^ for mESC, Bartsch *et al* Sci. Adv. 2023^46^ for hESC, and Susanto *et al*. Mol. Cell 2024^47^ for E12.5 mouse forebrain. **(F)** PANTHER protein classification analyses of proteins found in at least 3 rRNA IP replicates but absent in control IP samples, excluding core ribosomal proteins, translation factors, and known RAPs, reveal significant overrepresentation of proteins involved in RNA metabolism and translation for both subtypes.

To prepare biochemical input from labeled and purified CPN and SCPN for ribosome pulldown, we dissected labeled neocortices, dissociated labeled neurons for FACS purification, immediately lysed FACS-purified neurons with detergent, and performed centrifugation to remove nuclei and debris. To ensure rigorous control of rRNA IP-MS sample quality, we collected enough somata for both rRNA IP and control IP and quality control (QC) samples to monitor each step of the protocol (Figure 1C-D). Based on previous observations that SCPN somata are approximately twice the volume of CPN somata (∼1.25-1.3X diameter), we immunoprecipitated ribosomes from 50,000 SCPN or 100,000 CPN for MS samples to achieve comparable input (Table S1), in addition to quantitative normalization of input during later bioinformatic analysis.

To recover endogenous RCs from these very low-input samples, we developed an affinity purification approach using a monoclonal antibody against ribosomal RNAs (clone Y10B^37,38^). This antibody has been used to for immunocytochemical detection of ribosomes and verification of interactions between ribosomes and candidate associated proteins in several neurobiological studies^39–42^. During pilot studies on FACS-purified CPN, we initially tested an alternative approach with Cre-inducible HA-tagged RPL22/eL22 (RiboTag)^43^, which is commonly used to capture ribosome-associated transcripts for cell type-specific analysis from bulk tissue and has previously used as a positive control for affinity pulldown of ribosomes^23,25^. In the RiboTag approach, we used Emx1^IRES-Cre^ to induce expression of HA-tagged RPL22/eL22 in all cortical PN, and performed HA pulldown on FACS-purified retrograde-labeled CPN. We found that the Y10B-based rRNA IP method recovers ∼3 times the amount of 28S and 18S rRNA compared to the RiboTag HA IP method with FACS-purified CPN (Figure S1). Additionally considering that RPL22/eL22 is known to have extra-ribosomal functions and pull down proteins independent from assembled ribosomes^25,44^, we opted for the rRNA IP approach for these investigations. We employ retrograde labeling and FACS to obtain strict subtype specificity.

Further, based on QC samples that were split from samples prepared for MS, the Y10B-based rRNA IP approach shows high efficiency in pulling down ribosomes, recovering most of the available 18S and 28S rRNA from both purified CPN and SCPN (Figure 1C-D). In contrast, there is nearly undetectable yield from control IP samples using an isotype control antibody, indicating the stringency of washing steps in this IP protocol (Figure 1C-D).

We employed liquid chromatography with tandem mass spectrometry (LC-MS/MS) to investigate the proteins recovered with rRNA IP from purified SCPN and CPN. Based on calculations from QC samples, we selected rRNA IP samples with at least 100ng of 18S and 28S rRNA, and their control samples, for MS (Table S1). We used label-free quantification (LFQ), wherein samples are assayed sequentially, not simultaneously, aiming to most rigorously establish which proteins are truly present in some samples but absent in others. This capability to determine absence vs. presence of MS-detectable peptides distinguishes LFQ from approaches that assay samples simultaneously such as isobaric tandem mass tag (TMT) labeling, which standardly offers advantages in relative quantification studies. Advances in sample preparation, instrumentation, and data analytical algorithms now enable application of LFQ in quantitative differential analysis, even in the ultra-low-input regime^45^. We applied two forms of normalization to ensure rigor of identification of differential proteins (discussed below).

Proteomic analysis confirms that this rRNA IP approach enables access to the ribosome-associated complexes from both purified SCPN and CPN (Figure 1E-F, Table S2). We observe striking differences between rRNA IP samples (5 SCPN and 4 CPN replicates) and control IP samples (5 SCPN and 3 CPN replicates). We identify 76-77 of ∼80 known RPs from each rRNA IP sample for both subtypes. In contrast, we identify at most 11 RPs among all control samples combined, with several samples showing no identification of RPs whatsoever. Notably, we did not identify structural proteins of mitochondrial ribosomes in any rRNA IP samples, indicating that our rRNA IP approach specifically targets cytoplasmic ribosomes. Additionally, we identify between 200 to 300 co-immunoprecipitated proteins from each rRNA IP sample. These proteins include: 1) initiation and elongation translation factors; 2) proteins belonging to stringently filtered sets of RAPs previously reported in murine embryonic stem cells (mESC^25^) or human embryonic stem cells (hESC^46^); and 3) additional putative RAPs identified in bulk mouse forebrain at embryonic day (E) 12.5^47^. Further, protein classification analyses of proteins not classified as either RP, translation factors, or RAPs produced significantly enriched protein classes involved in RNA metabolism and translation (Figure 1F). The specific identification of most cytoplasmic RPs as well as translation factors, known RAPs, and RNA-processing and translation-related proteins confirms that this experimental approach successfully recovers ribosomal complexes from both SCPN and CPN *in vivo*. We focus here on comparative, differential analysis of proteins identified between SCPN and CPN. While some potential non-specific binding of non-RC proteins to Y10B cannot be entirely excluded, our analysis subtracts out non-specifically bound proteins, expected to be-common to both subtypes.

### SCPN and CPN ribosomal complexes share similar RP composition, but interact with distinct associated proteins

To address whether SCPN and CPN might have distinct subtype-specific ribosomal complexes, we directly compared rRNA IP-MS samples from SCPN with those from CPN, performing two levels of comparative analyses. First, we compared the composition of proteins present in SCPN and CPN ribosomal complexes, aiming to identify proteins exclusively detected in only one subtype. We analyzed the MS output of all rRNA IP samples grouped together, using Proteome Discoverer 3.2.’s “Label-free quantification” workflow. This analysis checks whether any peptide directly identified (via its MS2 spectra) in any rRNA IP sample has detectable abundance (MS1 peak) in other samples (Table S3). Second, among proteins detected in both subtypes, we performed differential analysis to identify proteins that might be significantly enriched in ribosomes of either subtype (Figure 2A, D).

**Figure 2.**
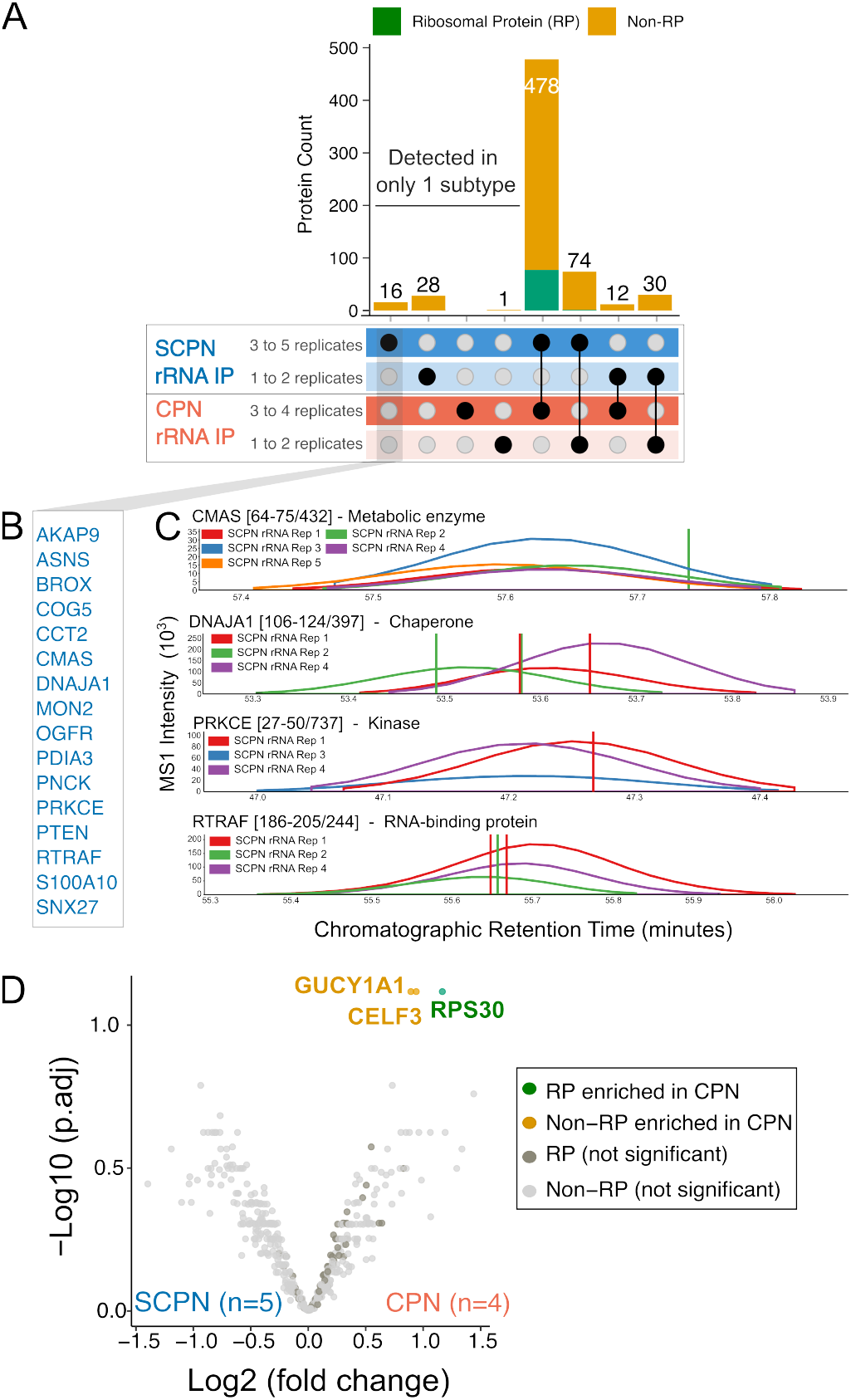
SCPN and CPN ribosomal complexes share similar RP composition, but interact with distinct associated proteins. **(A)** Label-free quantification proteomic analysis comparing SCPN and CPN ribosomal complexes: All identified RPs have detectable abundance in ribosomes of both sub-types. 16 associated proteins are detected exclusively in 3 or more replicates from SCPN. **(B)** List of 16 proteins detected exclusively in 3 or more replicates of rRNA IP from SCPN **(C)** Examples of extracted ion chromatograms of peptides from proteins that are detected exclusively in SCPN rRNA IP, showing only samples with detected abundance of indicated peptides. Peptides are referred to by their parent proteins and their amino acid sequence locations within their protein. Vertical lines indicate retention times of MS2 spectra used for peptide identification, with colors corresponding to sample origins of the MS2 spectra. A set of proteins with distinct known functions is shown. **(D)** Quantitative differential analysis of 478 proteins shared by at least 3 replicates of both subtypes: SCPN and CPN ribosomes share largely the same RP composition; 3 proteins enriched in CPN ribosomal complexes (adjusted p-value <=0.1)

Our analyses reveal key differences in the composition of non-core proteins between RCs of SCPN and CPN. We identify 16 proteins that are reproducibly detected in 3 to 5 replicates of SCPN rRNA IP samples, but not in any replicates of CPN rRNA IP samples (Figure 2A, Table S4). We personally inspected the extracted ion chromatograms of peptides belonging to proteins detected in only SCPN samples but not in any CPN samples, and confirmed in each case that only SCPN samples have detected peptides of these proteins. A few extracted ion chromatogram examples are shown in Figure 2C. Intriguingly, these 16 proteins have diverse known molecular functions, including metabolic enzymes, chaperones, kinases, and an RNA-binding protein (Figure 2C). In contrast to associated proteins, we find that all 78 identified core RPs are shared by RCs of both SCPN and CPN. 77 RPs belong to a set of 478 proteins that are reproducibly detected in at least 3 replicates of both subtypes (Figure 2A, Table S4).

We next sought to identify proteins quantitatively enriched in RCs of one or the other specific subtype, among the 478 shared proteins. Our bioinformatic workflow addresses both potential differences in input and technical variations among samples via two normalization approaches, and imputes any remaining missing 1-2 values for each subtype group (see Methods), thus enabling quantitative comparison (Figure S2). Our primary normalization approach, employing “median-of-ratios” normalization across all detected proteins for robustness against outliers and technical variability, results in similar overall distributions of protein abundances across CPN and SCPN samples (Figure S2A). This provides confidence in quantitative comparison between CPN and SCPN samples in the ultra-low-input regime. Subsequent differential analysis identifies three proteins that are significantly enriched in RCs of CPN (Figure 2D, Table S5). Among these proteins, RPS30/eS30 is the sole core ribosomal protein (it is a core component of mature 80S ribosomes, and it associates with immature ribosomes^48^), while CELF3 is an RNA-binding protein, and GUCY1A1 is a component of guanylate cyclase. Since RPS30/eS30 is produced by cleavage of a fusion protein comprising ubiquitin-like FUBI and RPS30/eS30^49^, we confirmed that the peptides used for RPS30/eS30 identification appropriately map to only the amino acid sequence of RPS30/eS30. None of the other RPs exhibit subtype-specific enrichment in RCs. Employing a second normalization approach, rescaling each sample to the average intensity of its core ribosomal proteins alone, differential analysis reveals equivalent CPN > SCPN enrichment of RPS30/eS30, GUCY1A1, and CELF3 (Figure S2 B, C). These three proteins rank among the five proteins with lowest p-values, though false-discovery-rate-corrected significance is reduced. Taken together, results from both protein detection analysis and quantitative analysis indicate that CPN and SCPN have distinct RCs.

### Proteins exclusive to SCPN rRNA IP samples physically interact with ribosomal components

To independently validate differences in proteins between SCPN and CPN ribosomes, we used two complementary approaches: verifying the physical interaction with RCs of proteins with differential abundance in SCPN vs CPN rRNA IP samples; and investigating the expression of these proteins in SCPN and CPN. We focused these investigations on the 16 noncore proteins reproducibly and exclusively detected in SCPN rRNA IP samples, since these likely represent the most strikingly divergent proteins between RCs of the two subtypes.

Among these 16 proteins, 11 proteins have previous independently reported evidence of physical interaction with core RPs (Figure 3A). We compiled known interactions between the 16 candidates and core RPs using BioGrid and OpenCell. BioGrid aggregates interactions discovered via multiple methodologies of interaction proteomics^50^; we restricted our search to the most stringent evidence using affinity purification, co-fractionation, and cross-linking experiments. OpenCell maps endogenous protein-protein interactions in human cell lines, using CRISPR editing of epitope tags for affinity purification and mass spectrometry^51^. Several candidates also interact with each other (Figure 3A).

**Figure 3.**
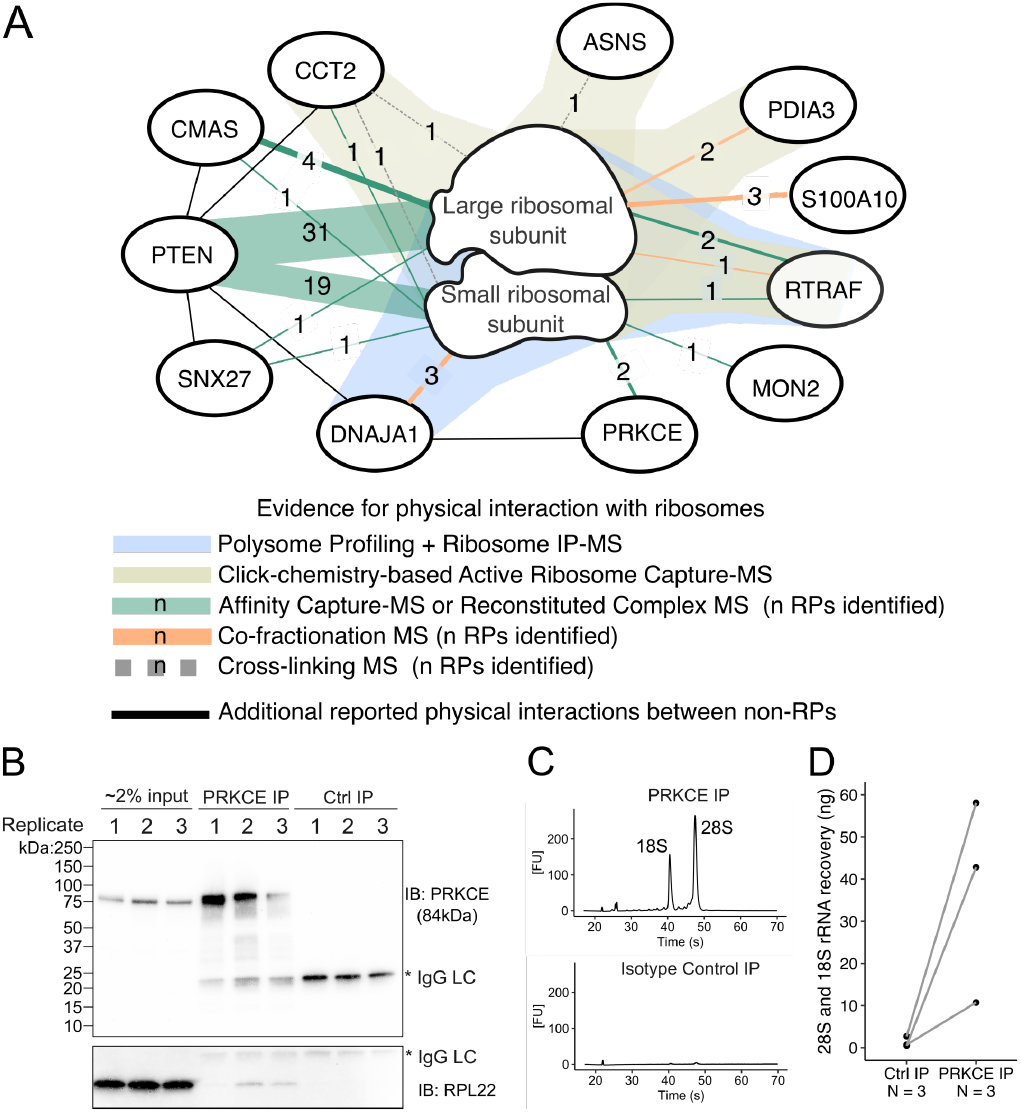
Ribosome interaction for proteins identified exclusively in SCPN by rRNA pulldown, with validation of PRKCE-ribosome interaction. **(A)** Protein interaction network showing reported physical interactions of SCPN-specific RAPs with ribosomal components and with each other. Line colors and thicknesses indicate distinct methodologies and number of interactions with ribosomal proteins (RPs) identified, respectively. Data were retrieved from BioGrid^50^, OpenCell^51^, Simsek *et al*. Cell 2017^25^, and Bartsch *et al* Sci. Adv. 2023^46^. **(B)** Co-immunoprecipitation of PRKCE and RPL22 (n = 3, biological replicates) **(C)** Specific recovery of ribosomal RNA in PRKCE IP versus paired isotype control IP, shown by representative Bioanalyzer electropherograms. **(D)** Quantification of 28S and 18S rRNA recovery following PRKCE IP versus paired isotype control IP.

Several of these 16 proteins also have previously reported evidence of interactions with ribosomes. Both DNAJA1 and RTRAF have been identified via mass spectrometry analysis of ribosomes recovered by a combination of polysome profiling and ribosome pulldown in mESC^25^. Their interactions are resistant to RNAse and puromycin, indicating that these proteins are *bona fide* ribosome-associated proteins, which interact with ribosomes independent of mRNA or nascent polypeptides^25^. RTRAF, along with CCT2, PDIA3, and ASNS have also been identified as RAPs by an orthogonal approach applied in hESC, which employs click chemistry on nascent peptides to isolate actively translating RCs, followed by biochemical elution of RC proteins (excluding nascent peptides) and MS^46^. A recent cryo-electron tomography study of the ribosome-ER translocon-oligosaccharyltransferase complex also suggests through structural prediction that PDIA3, an ER-resident chaperone for glycoproteins, matches an unassigned density in the structure^52^.

Intriguingly, these identified SCPN-specific ribosome-associated proteins with known interactions with components of ribosomes have diverse known functions. Both CCT2 and DNAJA1 are chaperones, in addition to ER-lumen chaperone PDIA3. Other proteins include: kinases PTEN and PRKCE; regulators of trafficking across compartments, including endosomal trafficker SNX27 and endosome-to-Golgi trafficking regulator MON2; and RNA-binding protein RTRAF. Meanwhile, both ASNS and CMAS are metabolic enzymes: ASNS catalyzes the conversion of aspartate and glutamine to asparagine and glutamate, respectively, while CMAS synthesizes cytidine 5-prime-monophosphate N-acetylneuraminic acid.

We further validated PRKCE for association with RCs in the developing mouse cortex. PRKCE is particularly intriguing because of its regulatory role in synaptogenesis; it inhibits dendritic spine development in immature neurons^53^, but promotes synapse formation and maturation in mature neurons, through phosphorylation of multiple substrates^54^. Incubating cytoplasmic lysate of P3 brain homogenate with a monoclonal antibody against PRKCE, we confirm co-immunoprecipitation both of ribosomal protein L22 and of 28S and 18S rRNA (Figure 3B-D). Thus, while BioGrid reports that PRKCE has previously known interaction with two proteins of the small ribosomal subunit (RACK1 and RPS27A)^50^, our results from these experiments using optimized buffer for ribosome stability further confirm that PRKCE associates with components of both ribosomal subunits.

### Several SCPN-specific, validated RC-associated proteins exhibit transcriptional enrichment and concordantly more translation in SCPN

We focused investigation on proteins both detected exclusively in SCPN RCs and with independent evidence of interaction with components of ribosomes. We investigated whether they show differential gene expression by SCPN versus CPN, potentially underlying their subtype-specific incorporation into RCs. We integrated our recently reported comparative transcriptomic and translational analysis of purified SCPN and CPN somata at P3^21^. We identify a subset of validated SCPN-specific RC-associated proteins with transcriptional enrichment in SCPN compared to CPN: S100A10, PRKCE, CMAS, ASNS, DNAJA1, PDIA3, and CCT2 (Figure 4A). For most of these proteins, their subtype-specific expression is maintained at the translational level (Figure 4B). Specifically, we find significantly more ribosome-protected fragments of S100a10, Asns, Prkce, Cmas, and Dnaja1, indicating higher translation of these transcripts by SCPN compared to CPN. For S100a10, our lab’s previously reported microarray data across development, and *in situ* hybridization data from P3 brain tissue, also reveal strikingly restricted S100a10 expression in SCPN in cortical layer V^2^ (Figure 4C).

**Figure 4.**
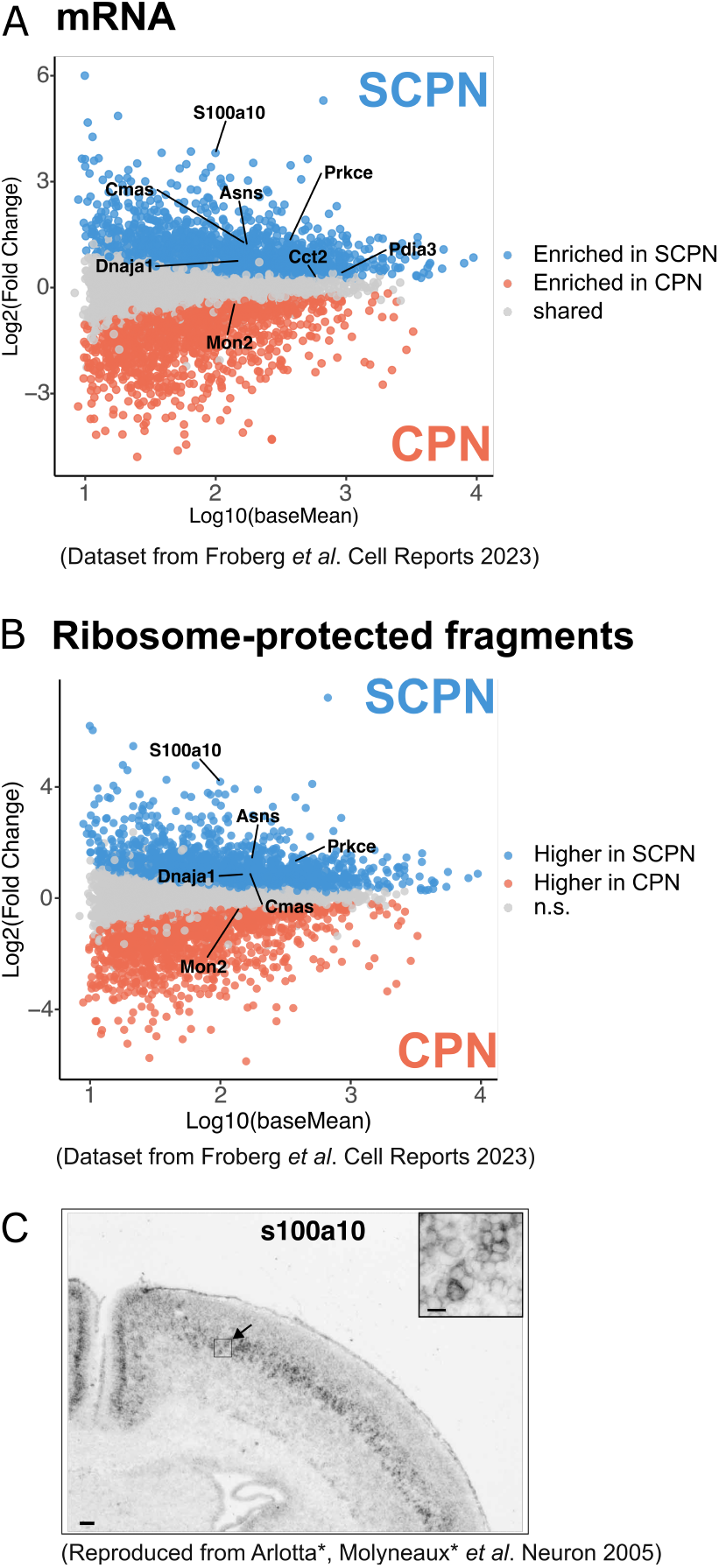
Several proteins both identified exclusively in RCs of SCPN and with validated ribosome interaction exhibit enriched expression and concordant translation by SCPN. **(A)** Comparative transcriptomic analysis of purified SCPN and CPN somata, highlighting differentially enriched transcripts encoding validated SCPN-specific RAPs. Dataset from Froberg *et al., Cell Reports* 2023^21^ **(B)** Comparative ribosome profiling of purified SCPN and CPN somata, highlighting transcripts encoding validated SCPN-specific RAPs with differential ribosome occupancy. Dataset from Froberg *et al., Cell Reports* 2023^21^ *(C) S100a10 ISH signal* is limited to cortical layer V, the predominant laminar position of SCPN. Reproduced from Arlotta*, Molyneaux* *et al. Neuron* 2005^2^, also reporting SCPN-specific expression compared with CPN.

These expression and translation differences between SCPN and CPN strongly suggest that this group of identified SCPN-specific ribosome-associated proteins have their origin as differentially associated with SCPN ribosomes via their subtype-specific expression. The others, which are not differentially expressed, might be differentially associated with SCPN ribosomes due to their post-translational processing, selective association with SCPN-specific proteins, or other factors.

## Discussion

In this report, we investigated whether ribosomal complexes from two distinct subtypes of cortical projection neurons (subcerebral output projection neurons and interhemispheric, associative projection neurons) might differ in protein composition during development while each subtype is executing long-range axonal extension and targeting. SCPN and CPN both arise from related dorsal “pallial” cortical progenitors in essentially the same cellular environment, but they extend their axons along strikingly distinct axonal trajectories and establish equivalently distinct circuitry, leading to quite distinct functions^1,7^. Translational control plays critical roles in neuronal differentiation and axon development^8,16^, but how such regulation is specialized by distinct neuronal subtypes during precise axonal targeting remains poorly understood. Ribosomes, rather than being fixed and homogeneous translational machinery, exhibit variations in their core protein composition, and interact with hundreds of associated proteins^22,23,25,55^, providing strikingly broader potential for diversity of cell type- and subtype-specific ribosome-associated complexes. There are increasingly more reported examples of specific core and associated proteins favoring or repressing translation of specific gene subsets^23–25,27,28^. Investigation of ribosomal complexes from distinct neuronal subtypes might enable new understanding of how regulation of translation contributes to dynamic subtype-specific circuit formation.

We developed an approach combining subtype-specific neuronal retrograde labeling, FACS purification of subtype-specific somata, ribosomal RNA pulldown, and ultra-low input mass-spectrometry to investigate the potential of subtype-specific RCs. Our approach offers several advantages. First, retrograde labeling definitively identifies subtype-specific neurons based on their axonal connectivity, thus is readily adaptable to diverse neuronal populations without the need for specific genetic drivers (often combinatorial and often non-specific at subtype level). Second, by isolating endogenous RCs from as few as 50,000 FACS-purified native neuronal somata, we both minimize potential functional perturbations from epitope tagging while preventing contamination from non-target cells—a significant issue when performing pulldowns from bulk tissue containing mixed labeled and unlabeled populations. Third, this approach pulls down ribosomal RNA rather than specific core ribosomal proteins, making it agnostic to the protein composition of potentially diverse ribosomes.

SCPN and CPN employ RCs with distinct protein compositions. Intriguingly, most of subtype-specific differences emerge from associated, non-core proteins. We identify 16 associated proteins enriched in RCs of SCPN to the point of not being detectable in CPN samples, and 2 associated proteins detected in both subtypes but quantitatively enriched in CPN samples. Focusing on SCPN-specific associated proteins, we identify 11 with previously reported interaction with core ribosomal proteins, and we further validate interaction with ribosomes by PRKCE. A subset of these proteins, including PRKCE, also exhibit enriched gene expression and concordantly more translation by SCPN compared to CPN, strongly suggesting that their subtype-specific association with RCs is established via subtype-specific expression. For other proteins without enriched expression by SCPN, their subtype-specific association with ribosomes might be regulated through post-translational modifications, interaction partners, subcellular localization, and/or other processes.

We note with interest that core RPS30/eS30 is enriched in RCs of CPN, strongly suggesting at least limited neuronal subtype-specific heterogeneity in 80S ribosomes. RPS30/eS30 resides at the solvent-exposed surface of ribosomes and exhibits the most dynamic exchange rate among RPs in mature ribosomes^56^, though its function in mature ribosomes and translation is essentially uninvestigated. Ribosome heterogeneity during cortical development is also supported by previous work using bulk cortex, with an increase in the level of RPL7/uL30 and a decrease in the level of RPL10/uL16 in polysomes between embryonic day (E) 13 and P0^13^. These shifts correspond to shifts in polysome-associated transcripts, and these changes are dependent on WNT3 produced from thalamic projections into the cortex^13^. Together, this prior study and our results strongly suggest that core ribosome composition might be regulated along multiple axes of subtype identity and developmental stage. Future mechanistic studies offer potential to elucidate roles of RPS30/eS30, e.g. in selective transcript recruitment to ribosomes, which would contribute to understanding of ribosome heterogeneity and its function in circuit formation.

The work presented here provides a complementary but distinct perspective to our lab’s parallel work quantifying transcript-specific translation by SCPN and CPN during the same range of developmental time^21^. nanoRibo-seq reveals that, while translational output largely follows transcript abundance, approximately 40 transcripts are an exception, with substantial subtype-specific differences in their translational efficiencies. In our current study, we identify a diverse set of proteins with subtype-specific association within heterogeneous ribosomal complexes. These proteins have known roles as RNA-binding proteins, chaperones, protein trafficking proteins, metabolic enzymes, and kinases. The diversity in functions of these associated proteins aligns with recent findings regarding ribosome-associated proteins more broadly, identifying proteins beyond the expected translation factors and known RNA-processing proteins^25,47^. Together, these results not only suggest potential regulators of the differentially translated transcripts identified by nanoRibo-seq, but also point to regulatory mechanisms beyond those directly affecting translational output and efficiency, e.g. those that might instead act on several important co-translational processes. We discuss several such proteins below.

Among the subtype-specific associated proteins, two exhibit RNA-binding capabilities that might enable regulation of transcript-specific translation: CELF3, enriched in CPN ribosomes, and RTRAF, specific for SCPN ribosomes. CELF3, a member of the CUG-binding protein and ETR-3-like factors (CELF) family, is an RNA-binding protein that functions in alternative splicing and is predominantly expressed in brain^57,58^. The CELF3 homolog in *Xenopus* has been shown to enhance translation of specific mRNAs, though such function has not been confirmed in mammals^59^. We did not identify enrichment of CELF3 binding motifs among the 40 transcripts that are differentially translated between CPN and SCPN, perhaps due to the small list of transcripts. In contrast, RTRAF functions in shuttling RNA between the nucleus and cytoplasm^60^, participates in RNA-transporting granules in neurons^61^, and forms a translation-initiating cap-binding complex^60^. However, its binding motifs remain unknown, precluding motif enrichment tests among the differentially translated genes. Additional work beyond the scope of this manuscript might focus on perturbation of CELF3 and/or RTRAF in one subtype, assessing potential impact of such perturbation on translational output within that subtype.

Focusing on proteins exhibiting subtype-specific association with RCs of SCPN, we identify proteins with diverse functions beyond RNA-binding, which suggests additional mechanisms for differential translational control. Several are chaperones, including CCT2, DNAJA1, and PDIA3, which suggests subtype-specific regulation of protein folding during translation. In addition, since PDIA3 is an ER-lumen chaperone with likely association with the ribosome-ER translocon complex^52^, its detection exclusively in SCPN suggests that SCPN might have a higher proportion of ER-localized ribosomes compared to CPN.

In contrast, while PRKCE exhibits both validated physical interaction with ribosomal complex proteins and SCPN-specific gene expression, its role as a kinase with diverse substrates does not immediately suggest clear mechanistic connections to ribosome regulation and translational control. While PRKCE pulldown from mouse heart tissue recovers numerous RNA-binding proteins, these interactions might be mediated through ribosomal complexes^62^. Nevertheless, it is intriguing that it inhibits synaptogenesis in immature neurons while promoting formation of synapses by mature neurons through phosphorylation of diverse substrates^53^.

It remains possible that the observed subtype-specific differences in RCs might be related to a broader set of biological processes beyond axonal connectivity. For example, at the time of purification at P3, SCPN and CPN might also be acquiring other aspects of their respective identities, including morphology, dendritic connectivity, etc.. In addition, since SCPN are born around E13.5 while CPN production spans between E13.5 and E17.5, SCPN and CPN might also be implementing different stage-specific processes at the time of soma purification at P3. RCs might thus be specialized for specific stage and subtype-specific needs of PN development. Our work presented here provides a starting point for further investigation of such RC specialization, as well as investigation of potential RC specliazation corresponding to further diversity and heterogeneity within both CPN and SCPN subtypes.

A few future directions might further elucidate how proteins comprising subtype-specific RCs might regulate translational control and circuit formation. First, it would be highly informative to further validate and characterize the nature of the interactions between associated proteins and ribosomes. Such investigations could be conducted in subtype-specific somata by building on our existing approach, with: 1) addition of RNase treatment to determine whether a protein-ribosome interaction is mRNA dependent; and/or 2) addition of a range of ribosome inhibitors to capture distinct functional states. Regarding approaches orthogonal to immunoprecipitation, sucrose gradient fractionation for polysome profiling is likely infeasible by currently available methods, given the limited material of subtype-specific PNs. Imaging approaches, including proximity ligation assays and super-resolution microscopy, are likely more applicable but would require extensive candidate-specific protocol optimization. Second, manipulation of expression of select proteins in specific circuitry (for example, by *in utero* electroporation of plasmids in mice at embryonic day 13.5 or 15.5 to manipulate gene expression in cortical layer V neurons (including SCPN) or layer II/III CPN, respectively) would enable functional investigation, particularly with regard to function in establishment of precise connectivity and circuitry. Third, TMT labeling with alternative approaches to normalization could be employed to better define relative quantitative differences between functional candidates present in multiple subtypes. Fourth, extension of this work to subcellular axon and growth cone investigation, though currently not feasible due to exceptionally low available input, would enable elucidation of critical local translational regulation and direct comparison with regulation within somata of the same neuron subtypes.

Together, these results indicate that ribosomal complexes exhibit subtype-specific protein composition in distinct subtypes of cortical projection neurons during development, and identify both potential candidates for further investigation and a potential range of functions they might likely exert in translational regulation critical for precise and diverse subtype-specific circuit formation and function.

## Acknowledgements

We thank Dr. Y. Itoh of our lab for the generation of Emx1^IRES-Cre^Rpl22-3xHA^fl/fl^TdTomato^fl/fl^ mice; J. Heo, J. Iyer, M. Ross, B. Wall, and G. Wheeler for technical assistance; D.E.Tillman for expertise and analysis and S. Morton for support with proteomic data analysis; Dr. K. Ozkan for schematics; other members of our laboratory for scientific discussions and helpful suggestions; Dr. M. Barna and members of the Barna lab at Stanford for insightful discussions and suggestions on a few approaches for ribosome pulldown; J. LaVecchio, N. Kheradmand, and G. Kassis of the HSCRB-HSCI Flow Cytometry Core; and the Harvard Center for Biological Imaging (RRID:SCR_018673) for infrastructure and support. This work was supported by grants to J.D.M.: National Institutes of Health (NIH) Pioneer Award DP1 OD/NS106665 and RF1 AG083085, and the Max and Anne Wien Professor of Life Sciences fund, with additional infrastructure support by NIH NS045523 and NS104055. J.E.F. was partially supported by NIH T32 AG000222, and NIH F32 AG067661. This paper was typeset with the bio-Rxiv word template by @Chrelli: www.github.com/chrelli/bioRxiv-word-template.

## Author contributions

Research designed by T.P.T. and J.D.M.; research performed by T.P.T. and B.B; data analyzed by T.P.T., B.B, J.D.M; and manuscript written by T.P.T. and J.D.M, with input from the other authors.

## Materials and Methods

### Mice

The experimental procedures using mice were reviewed and approved by the Harvard University Institutional Animal Care and Use Committee (protocol number HU IACUC ID # 11-22), and were performed following institutional and federal guidelines. We ordered timed pregnant CD-1 (RRID:IMSR_CRL:022) dams for all experiments, except for those with RPL22-3xHA^fl^ (RiboTag) mice. For pilot experiments with RiboTag mice, we crossed Emx1^IRES-Cre^ (JAX stock #005628^63^), Rpl22-3xHA^fl^ (JAX #011029^43^), and Ai9(RCL-tdT or TdTomato ^fl^) strains (JAX stock #007909^64^) to generate Emx1^IRES-Cre^ RPL22-3xHA^fl^ TdTomato^fl^ mice. A recombination event enabled the Emx1^IRES-Cre^ and TdTomato^fl^ alleles to be on the same chromosome 6. The cross between RPL22-3xHA^fl/fl^ TdTomato^fl/fl^ male and Emx1^IRES-Cre/+^ RPL22-3xHA^fl/fl^ TdTomato^fl/fl^ female mice produced both Cre-induced mice and “no Cre” littermate controls for experiments. For retro-grade labeling, we performed ultrasound-guided injections on P1 littermates, targeting the corpus callosum to label CPN, or the cerebral peduncle to label SCPN. All experiments used mixed groups of male and female mice. We report developmental stages of all experiments in the descriptions of the experiments.

### Neuronal subtype retrograde labeling, dissociation and purification

We performed retrograde labeling of CPN and SCPN via ultrasoundguided injection of fluorophore-conjugated cholera toxin B (CTB) at P1-2, as previously described^2,65^. Briefly, we placed pups on ice for 3 mins to deeply anesthetize them by hypothermia, then secured them gently on the injection platform. We used ultrasound backscatter microscopy to visualize injection sites and to guide the injection micropipette. For CPN labeling, CTB was injected into the right side of the corpus callosum, with four to five injection sites along the rostral-caudal axis. For SCPN labeling, we injected at six sites within the left cerebral peduncle (two sites along the dorsal-ventral axis along each of three penetrations along the medial-lateral axis). Each injection site received five 25 nL pulses. Either AlexaFluor 488 or 647 were conjugated to CTB; the specific fluorophore is included in the descriptions of individual experiments.

We performed neuronal somata FACS essentially as described in Engmann and Hatch *et al*. Nat. Protoc. 2022^66^, with the modification that all buffers starting with enzymatic digestion (using papain and DNAse I) contained 100µg/mL cycloheximide (CHX) to stabilize the ribosomal holocomplexes. We dissected brains in ice-cold HBSS under epifluorescence, and placed microdissected tissue in dissociation solution (DS). We washed tissue twice in DS, then enzymatically digested twice by incubation in enzyme solution (DS with cysteine, papain, and DNAse I) and CHX. We washed twice in wash solution (WS) + CHX + 5mM MgCl_2_, and triturated 15–20 times in ∼1 mL of WS + CHX using fire-polished glass pipettes. We diluted cell suspensions with 4 mL WS + CHX + 5mM MgCl_2_, centrifuged to collect cells for 5 min at 80 × g-force, triturated again in 1 mL WS + CHX + 5mM MgCl_2_, and passed the cell suspension through a strainer cap. We added 1:1,000 SYTOX Blue to enable sorting of viable cells.

During FACS, we sorted SytoxBlue-negative and AF488-positive neurons, and collected them in 1.5mL Protein LoBind Eppendorf tubes. We included controls for all color channels to enable compensation and precise gating of fluorescence signal. The collection tubes were previously rinsed in 100% acetonitrile to remove contaminants, and were subsequently air-dried. Each collection tube contained 4 × polysome buffer, which consisted of 20mM HEPES, 150mM KCl, 20mM MgCl_2_, 400µg/mL CHX, 4 unit/mL RNasin® Plus Ribonuclease Inhibitor (Promega Cat# N2615), 2mM dithiothreitol (DTT), and 1 tablet/2.5mL cOmplete™, Mini, EDTA-free Protease Inhibitor Cocktail (Roche Cat# 11836170001). We used a volume of 4× polysome buffer equal to one third of the final volume of PBS used as sheath fluid for sorted somata (∼1.8nL PBS/somata).

We typically sorted 290,000 CPN or 145,000 SCPN per ribosome IP experiment. For SCPN, we diluted the final volume by 2-fold, using 3 parts PBS and 1 part 4 × polysome buffer, such that the final SCPN and CPN samples had the same volume. We added TurboDNAse (Invitrogen Cat# AM2238) to the final samples to reach a final concentration of 24U/mL, and added MgCl_2_ to reach a final concentration of 10mM.

### Endogenous ribosome pulldown in FACS-purified neurons

To prepare the cytoplasmic lysate from purified somata, we performed a rapid cellular lysis step by adding 0.25% Triton X-100 and mixing the lysate for ∼1 minute with end-to-end rotation. To remove the nuclei and cellular debris, we performed a rapid centrifugal spin at 4°C for 10 seconds until the speed reached 17,000 × g-force, then immediately performed a 5 min lower-speed centrifugation at 1,700 × g-force, before collecting the supernatant. We repeated this two-spin step and collected the resulting supernatant. Because the nucleus is the site of ribogenesis and early steps of maturation, our protocol removes a major source of immature ribosomes by subjecting FACS-purified cells to two centrifugal spins that remove the nucleus. In pilot experiments, nuclear removal was confirmed by the absence of a contaminating genomic DNA peak on Bioanalyzer electropherograms of total RNA extracted from input samples (without genomic DNA removal) immediately prior to rRNA-IP. We set aside 24µL (volume equivalent to the cytoplasmic lysate of 10k CPN somata or 5k SCPN somata) for RNA extraction with the RNeasy Plus Micro Kit (Qiagen Cat#74034) and to serve as a QC sample for input.

For endogenous ribosome pulldown, we split the sample into two identical 1.5mL tubes, each with the equivalent of the cytoplasmic lysate of 140k CPN somata or 70k SCPN somata. We used one tube for pulldown with Y10B antibody (Santa Cruz Biotechnology Cat# sc-33678, RRID:AB_628226), and the other for control pulldown with Mouse IgG2a, κ Isotype Ctrl Antibody (immunogen: keyhole limpet hemocyanin; BioLegend Cat# 401502, RRID:AB_2800437). We added 800ng of antibody per the equivalent of lysate of 100K CPN (or 50k SCPN somata). We incubated the samples at room temperature for 30 min, then at 4°C for 15min, with end-to-end rotation. To recover the antibody-ribosome complexes, we added 5µL of Protein G-conjugated magnetic beads (Thermo Fisher Cat#10003D) per 800ng of antibody, and incubated the samples for 1.5 hours at 4°C, with end-to-end rotation. We then used a magnetic stand to separate the magnetic beads from the flowthrough. We collected the flowthrough for RNA extraction using the the RNeasy Plus Micro Kit.

To remove potential non-specifically bound material from the magnetic beads-antibody complexes, we washed the magnetic beads four times with a high-salt wash buffer (20mM HEPES, 10mM MgCl_2_, 350mM KCl, 0.25% Triton X-100, 100µg/mL cycloheximide and 0.5mM DTT), with 1 minute of end-to-end rotation between the washes to resuspend the magnetic beads well in high-salt wash buffer. Before the last wash, we divided the magnetic bead suspension in wash buffer into two tubes: one contained the equivalent of the IP from the lysate of 100k CPN (or 50k SCPN somata), intended for snap freezing for MS, while the other tube contained the equivalent of the IP of the cytoplasmic lysate of 40k CPN somata (or 20k SCPN somata), intended for RNA extraction for QC of the IP. For both tubes, we rinsed the magnetic beads 3 times with PBS supplemented with 10mM MgCl_2_ and 100µg/mL CHX. We snap-froze the MS samples in liquid nitrogen. For the QC samples, we added 100µL of RNA Lysis Buffer from Zymo *Quick*-RNA Microprep Kit (Zymo Research Cat# R1050), briefly vortexed the magnetic bead suspension, and collected the liquid separated from the magnetic beads for downstream RNA purification using the standard Zymo *Quick*-RNA Microprep Kit protocol.

### Ribosome pulldown in FACS-purified CPN in RiboTag mouse

We prepared cytoplasmic lysates from purified somata as for endogenous ribosome pulldown. For HA pulldown, we used 5µL of Pierce™ Anti-HA Magnetic Beads (Thermo Fisher Cat# 88836) per cytoplasmic lysate of 100k CPN somata, performing parallel pulldown from samples with Emx1-Cre or without Cre. We incubated samples at room temperature for 30 min with end-to-end rotation. Subsequently, we performed the same wash protocol used with endogenous ribosome IP to remove non-specific binding from the magnetic beads (four times with high salt buffer and three times with PBS supplemented with 10mM MgCl_2_ and 100µg/mL CHX).

### Ribosomal RNA measurement

We used an Agilent 2100 Bioanalyzer system with the RNA 6000 Pico kit (Agilent), or an Agilent 4200 TapeStation system with the High Sensitivity RNA kit (Agilent) to analyze purified RNA samples from a range of QC samples collected during sequential steps of the pulldown. We quantified the concentration of 18S and 28S rRNA.

### Label-free quantitation mass spectrometry data acquisition

We submitted 19 samples (5 SCPN rRNA IP, 5 CPN rRNA IP, 5 SCPN control IP, and 4 CPN control IP) for ultra-sensitive mass spectrometry and proteomic analyses (Table S1). On-bead digestion was performed with 100 ng of trypsin for each sample, using a stock solution of Trypsin Platinum, Mass Spectrometry Grade (Promega Cat# VA9000 at 100ug/mL in 50mM TEAB. Samples were incubated for 2 hrs at 50°C and mixed at 350 rpm on an Eppendorf ThermoMixer. Digested samples were transferred to HPLC glass vials and resuspended in 6uL of 0.1% formic acid in ultrapure HPLC grade water, then injected for liquid chromatography and tandem mass spectrometry (LC-MS/MS).

Samples were analyzed sequentially on the same day, with each sample used for a single LC-MS/MS run. Control IP samples were analyzed before rRNA IP samples to avoid even minor potential sample carry-over from rRNA IP samples (no CPN Control IP Replicate 3, due to insufficient sorted somata for preparation) (Table S1). Within each IP type, all SCPN samples were analyzed consecutively before proceeding to CPN samples to minimize even minor potential cross-subtype contamination (Table S1). These LC-MS/MS runs were performed on an Orbitrap Exploris™ 240 Mass Spectrometer (Thermo Scientific) equipped with a NEO nanoHPLC pump (Thermo Scientific). Peptides were separated via a 75µm × 4 cm C18 trapping column (Premier LC, CA) followed by an analytical column PepMap Neo 50µm × 150mm (Thermo Scientific, Lithuania). Separation was achieved by applying a gradient of 5–25% acetonitrile in 0.1% formic acid over 60 min at 200 nl/min. Electrospray ionization was performed by applying a voltage of 2.1 kV using a PepSep electrode junction at the end of the analytical column, and sprayed from a stainless steel PepSep emitter SS with liquid junction of an inner diameter of 30µm (Bruker, MA). The Exploris Orbitrap was operated in the data-dependent mode for MS data acquisition. An initial MS survey scan was performed in the Orbitrap in the range of 450–900 mass/charge (m/z) at a resolution of 1.2 × 10^5^. Higherenergy collisional dissociation (HCD) MS2 scans were recorded after each MS1 scan for the ten most intense ions. For each HCD MS2 scan, the fragment ion isolation width was set at 0.8 m/z, automatic gain control (AGC) target at 50,000 ions, maximum ion injection time at 150 ms, normalized collision energy at 34V, and activation time at 1ms.

### Mass spectrometry data analysis

For peptide identification, the raw file from each control sample and rRNA IP sample was analyzed in Proteome Discoverer 3.2 software (Thermo Scientific, RRID:SCR_014477). Assignment of MS/MS spectra was performed with both Sequest HT and CHIMERYS™ (MSAID, Germany) algorithms, using a protein sequence database that included all entries from the Mouse Uniprot database (SwissProt 19,768 2019), and known contaminants (e.g. human keratins, other common lab contaminants). Sequest HT searches were performed using a 20 ppm precursor ion tolerance, requiring each peptide’s N- and C-termini to adhere with trypsin protease specificity, while allowing up to two missed cleavages. For these searches, carbamidomethylation of cysteine was set as a fixed modification, while variable modifications included oxidation of methionine and phosphorylation of serine, threonine, and/or tyrosine. An MS/MS spectra assignment false discovery rate (FDR) of 1% at the protein level was achieved by applying a target-decoy database search. Filtering was performed using Percolator (64 bit version^67^)

For analysis of protein detection between control and rRNA IP samples (Figure 1E), we retrieved protein-level data from individual searches and removed contaminant proteins and proteins without quantified abundance (Table S2). From these analyses, we identified a single QC failure, leading to sample exclusion: CPN control IP replicate 5 unexpectedly identified 362 proteins, strikingly more than all other control samples, which detected 6 to 19 proteins. Both the CPN control IP replicate 5 and CPN rRNA IP replicate 5, which came from the same input and were processed in parallel, were therefore excluded from subsequent analyses.

For Figure 1F, protein classification analyses were performed on proteins found in at least 3 rRNA IP replicates, but absent in all control IP samples, for each subtype, using PANTHER^68^. We used PANTHER Overrepresentation Test^69^ (released 20240807) with the following parameters: 1) background: mouse genome; 2) annotation data set: “PANTHER Protein Class”; 3) test type: “Fisher’s Exact”; and 4) correction: “Calculate False Discovery Rate”.

For comparative analysis of rRNA IP samples from CPN and SCPN (Figure 2), we additionally submitted the raw files of these samples as a group for analysis using the “label-free quantitation” workflow in Proteome Discoverer. This workflow performs chromatographic retention time alignment across samples and maps LC-MS (MS1) peaks in each sample with identified peptide-spectrum matches pooled from all samples. We then retrieved peptide-level ouput with per-sample peptide abundance (MS1 precursor ion intensity) (Table S3). Peptides without detected MS1 peaks were not assigned abundance values and considered not detected. To most stringently determine protein-level detection and quantification in each sample, we used only unique peptides, which map to single proteins and ensure unambiguous protein identification. In particular, protein-level abundances were calculated as the geometric means of all associated unique peptide abundances. Proteins for which there was no single unique peptide detected in a sample were not assigned abundance values for that sample. From an initial set of 651 proteins quantifiably detected via unique and/or non-unique peptides across all samples combined, 12 were removed that were detected solely from non-unique peptides (these proteins were detected in samples of both subtypes based on non-unique peptides). For each protein, we then counted the number of samples per subtype with detected abundance (Table S4). Notably, the 16 proteins detected in 3 or more replicates of SCPN samples but not in any CPN samples were detected via solely their unique peptides, eliminating the possibility of falsely missing their detection in CPN samples via non-unique peptides. Additionally, we manually inspected the extracted ion chromatograms of peptides belonging to these proteins, and confirmed that the detection of each protein is exclusive to SCPN.

For differential abundance analysis of proteins that are shared by both SCPN and CPN rRNA IP (Figure 2D, Table S5), we retained only proteins detected in 3 or more replicates of each of the CPN and SCPN groups. We applied two approachs to normalize the data. As a primary approach, “Median of ratios” normalization was applied to protein abundances to most rigorously account for differences in protein input across samples, in the ultra-low-input regime. Specifically, protein intensities were log-transformed, and a pseudo-reference was calculated as the geometric mean intensity of each protein across all samples. Then, for each sample, the difference of each protein’s log intensity to the pseudo-reference was calculated, and the median of these ratios was taken as the sample-specific scaling factor. Raw intensities were then divided by the exponentiated scaling factor to yield normalized values. We applied a second normalization, dividing protein abundances by the mean intensity of only ribosomal structural proteins in each sample, then subsequently cross-referenced results between the two approaches. Following normalization, to impute missing values in the remaining 1-2 samples of some subtype groups, we performed kNN imputation (k = 2) for CPN and SCPN samples separately. Protein abundances were then log2-transformed. Differential abundance analysis was conducted with Linear Models for Microarray (LIMMA), implemented by the DEP package^70^ from the Bioconductor repository (RRID:SCR_006442). Multiple testing correction was performed using the Benjamini-Hochberg method, and the cut-off for statistical significance was set at adjusted p-value <0.1 (Table S5).

All raw MS data and supplementary tables associated with MS data analysis are deposited in the Harvard Dataverse repository (https://doi.org/10.7910/DVN/ZQF9LQ). All code is available on GitHub at https://github.com/tienphuoctran/Subtype-specific-ribosomes_Macklis.

### Pulldown of candidate ribosome-associated proteins

To prepare IP input, we collected P3 cerebral cortex tissue from ∼3-4 pups/replicate in a Duall tissue grinder containing 1× polysome buffer (prepared by mixing 3 parts PBS and 1 part 4× Polysome buffer, as detailed in the method subsection for Ribosome IP from FACS-purified neurons). We used a ratio of 1g of wet tissue to 9.75 mL of 1× polysome buffer. We homogenized the tissue with 10-20 strokes on ice. We added Triton X-100 to a final concentration of 0.25% and mixed the samples with end-to-end rotation at 4°C for 10 minutes to enable detergent lysis. To collect the cytoplasmic lysate, we performed two centrifugal spins of 1,700 × g-force at 4°C for 15 min and collected the supernatant after each spin. For sample pre-clearing to remove components that might non-specifically bind to Protein G and magnetic beads, we incubated the sample with 10µL of Protein G-conjugated magnetic beads per 1 mL of lysate at 4°C for 45 minutes with mixing by end-to-end rotation. We collected the pre-cleared lysate by using a magnetic stand to separate the magnetic beads. Since we typically prepared 3 biological replicates at once, we normalized the replicates by total RNA concentration.

For immunoprecipitation, we prepared 2 identical 1.5mL tubes from each replicate, adding antibodies against a protein-of-interest into one and isotype control antibodies into the other. The antibodies and the amount used are as follows: PRKCE (BD Biosciences Cat# 610085, RRID:AB_397492, 2µg/1.5mL lysate) and Mouse IgG2a Isotype Control Antibody (BioLegend Cat# 401502, RRID:AB_2800437, 2µg/1.5mL lysate). We used the same amount of control antibody as we did for antibodies against candidate proteins. We incubated the samples at 4°C overnight with mixing by end-to-end rotation. We added 5µL of Protein G-conjugated magnetic beads per 800ng of antibody, and incubated the samples for 2 hours at 4°C, with end-to-end rotation. We used a magnetic stand to separate the magnetic beads from the flowthrough, and washed the magnetic beads three times with 1× polysome buffer (with 0.25% Triton X-100). To elute proteins for western blotting, we incubated the magnetic beads with 1× SDS sample buffer at room temperature for 5 min with constant mixing at 600 rpm before collecting the supernatant with the use of a magnetic stand. To extract total RNA, we added 100 µL of RNA Lysis Buffer from a Zymo *Quick*-RNA Microprep Kit, briefly vortexed the magnetic bead suspension, and collected the liquid separated from the magnetic beads for downstream RNA purification using the standard Zymo *Quick*-RNA Microprep Kit protocol.

### Transcardial perfusion, tissue processing, and microscopy

Postnatal mice were deeply anesthetized via hypothermia and then transcardially perfused with ∼5 ml of ice-cold PBS followed by 5 ml of ice-cold 4% paraformaldehyde. Brains were carefully extracted from the skull and post-fixed in 4% PFA at 4°C overnight with gentle mixing. On the following day, the brains were rinsed twice with ice-cold PBS and then placed in 30% (w/v) sucrose in PBS with gentle mixing for cryoprotection. Tissue was embedded in O.C.T (Sakura Finetek USA Inc Cat# 25608-930), frozen at -80°C for 30 minute then at -20°C for 2 hours before being sectioned using a cryostat (Leica, 50 µm thick coronal sections for immunocytochemistry). Sections were collected in PBS supplemented with 0.025% sodium azide.

For visualization of retrograde labeling using fluorophore-conjugated CTB (as in Figure 1B), sections were washed once with PBS, counterstained with DAPI (0.5µg/mL) for 10min, washed again three times with PBS, and then mounted on glass slides (Superfrost, VWR Cat#48311-702 VWR). We applied Fluoromount-G mounting media (VWR/Southern Biotech Cat# 0100-01) before placing the coverslip and sealing the slides with nail polish.

We acquired images of whole brain sections with an epifluorescence microscope equipped with a motorized stage and a 10x objective (Nikon NiE), using mosaic image stitching through NIS Elements software (Nikon). Images were processed using Fiji (RRID:SCR_002285)^71^.

## Supplementary Figures

**Figure S1.**
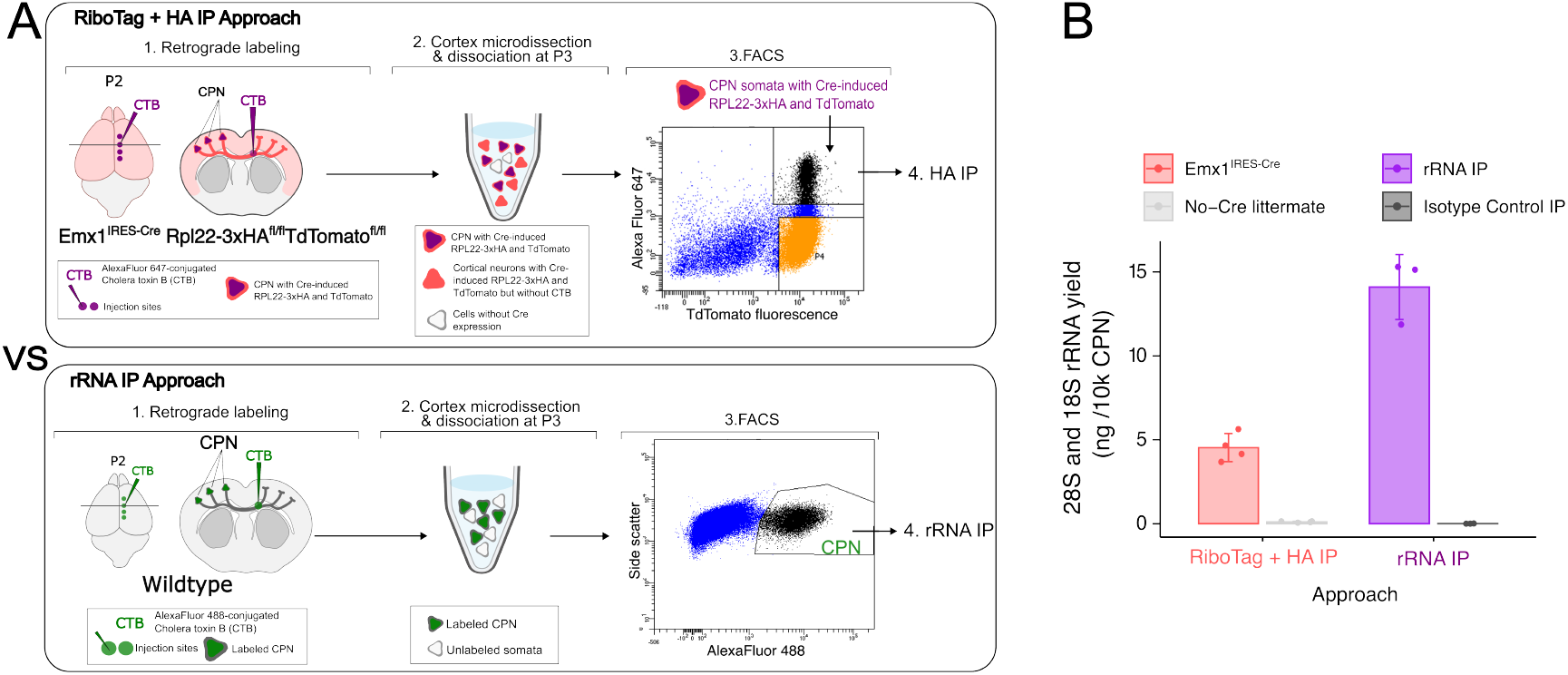
rRNA IP approach recovers more rRNAs from FACS-purified CPN than an approach using RiboTag mice and HA IP. **(A)** Schematic of pulldown of ribosomal complexes using HA IP from CPN expressing Cre-induced HA-tagged RPL22 (RiboTag + HA IP approach), compared to rRNA IP approach. **(B)** rRNA IP recovers ∼3-times the amount of 28S and 18S rRNA than HA IP from RiboTag mice, using the same number of FACS-purified CPN, while maintaining low non-specific binding. Individual dots represent biological replicates, with bars showing mean values and error bars indicating standard deviation.

**Figure S2.**
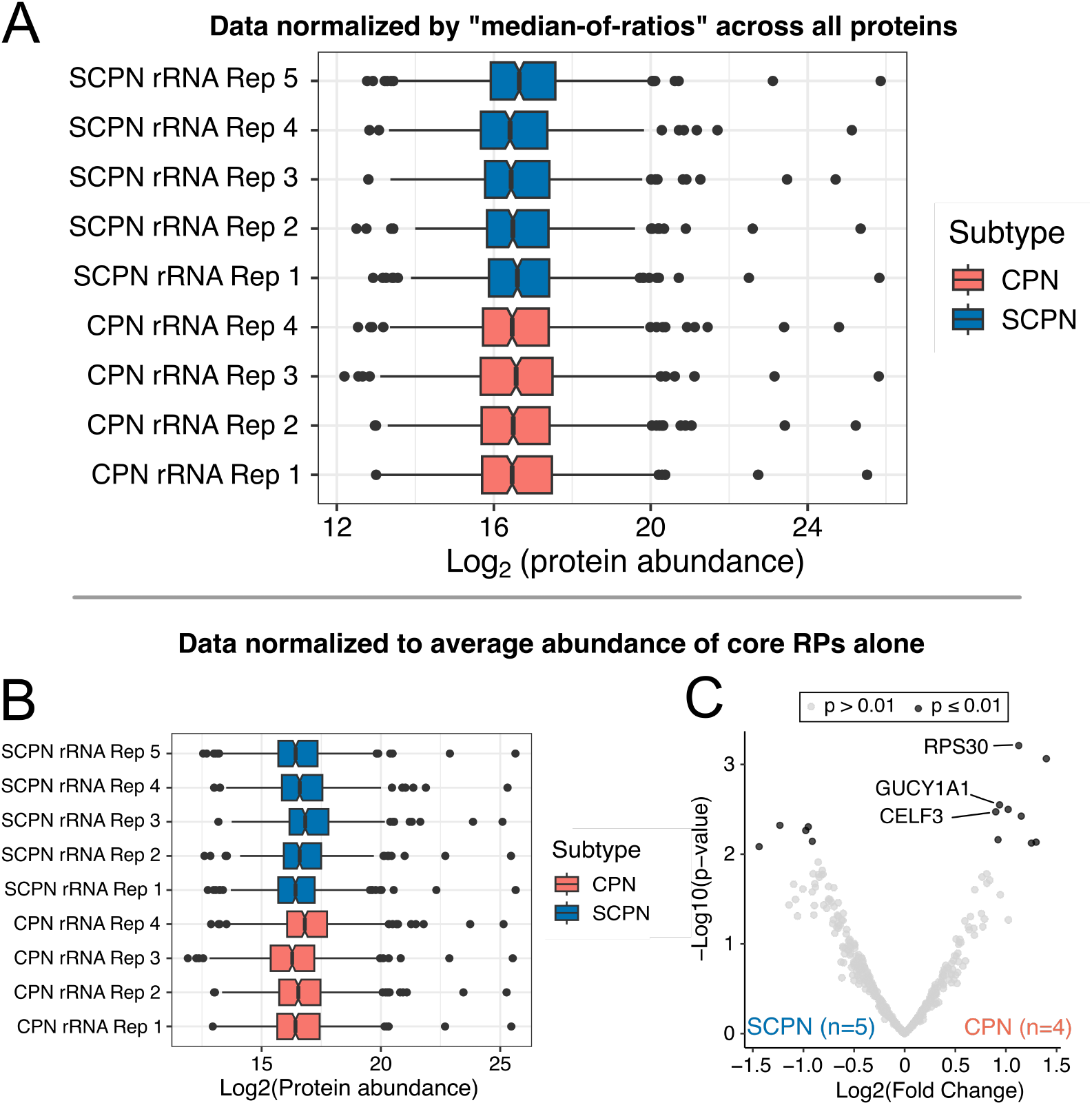
Bioinformatic normalization enables differential analysis between CPN and SCPN rRNA IPs replicate samples. **(A)** Median-of-ratios normalization was applied to protein abundances to account for differences in protein input between replicate (Rep) samples. For missing values in 1-2 samples within each subtype group, CPN and SCPN samples were analyzed separately using k-nearest neighbor (kNN) imputation (k=2). Protein abundances were then log2-transformed. **(B, C)** As a second, comparative normalization approach, protein abundances were normalized to the average abundance of core ribosomal proteins alone. Subsequently, the same kNN imputation and log2 transformation were performed. (C) This differential analysis reveals equivalent CPN > SCPN enrichment of the top candidates RPS30/eS30, GUCY1A1, and CELF3 identified in Figure 2.

## References

1. Greig, L.C., Woodworth, M.B., Galazo, M.J., Padmanabhan, H., and Macklis, J.D. (2013). Molecular logic of neocortical projection neuron specification, development and diversity. Nat Rev Neurosci 14, 755–769. 10.1038/nrn3586.

2. Arlotta, P., Molyneaux, B.J., Chen, J., Inoue, J., Kominami, R., and Macklis, J.D. (2005). Neuronal Subtype-Specific Genes that Control Corticospinal Motor Neuron Development In Vivo. Neuron 45, 207–221. 10.1016/j.neuron.2004.12.036.

3. Taylor, J.P., Brown, R.H., and Cleveland, D.W. (2016). Decoding ALS: from genes to mechanism. Nature 539, 197–206. 10.1038/nature20413.

4. Özdinler, P.H., Benn, S., Yamamoto, T.H., Güzel, M., Brown, R.H., and Macklis, J.D. (2011). Corticospinal Motor Neurons and Related Subcerebral Projection Neurons Undergo Early and Specific Neuro-degeneration in hSOD1G93A Transgenic ALS Mice. J. Neurosci. 31, 4166–4177. 10.1523/JNEUROSCI.4184-10.2011.

5. Cobos, I., and Seeley, W.W. (2015). Human von Economo Neurons Express Transcription Factors Associated with Layer V Subcerebral Projection Neurons. Cerebral Cortex 25, 213–220. 10.1093/cercor/bht219.

6. Lin, L.-C., Nana, A.L., Hepker, M., Hwang, J.-H.L., Gaus, S.E., Spina, S., Cosme, C.G., Gan, L., Grinberg, L.T., Geschwind, D.H., et al. (2019). Preferential tau aggregation in von Economo neurons and fork cells in frontotemporal lobar degeneration with specific MAPT variants. acta neuropathol commun 7, 159. 10.1186/s40478-019-0809-0.

7. Fame, R.M., MacDonald, J.L., and Macklis, J.D. (2011). Development, specification, and diversity of callosal projection neurons. Trends in Neurosciences 34, 41–50. 10.1016/j.tins.2010.10.002.

8. Zahr, S.K., Yang, G., Kazan, H., Borrett, M.J., Yuzwa, S.A., Voronova, A., Kaplan, D.R., and Miller, F.D. (2018). A Translational Repression Complex in Developing Mammalian Neural Stem Cells that Regulates Neuronal Specification. Neuron 97, 520–537.e6. 10.1016/j.neuron.2017.12.045.

9. Yang, G., Smibert, C.A., Kaplan, D.R., and Miller, F.D. (2014). An eIF4E1/4E-T Complex Determines the Genesis of Neurons from Precursors by Translationally Repressing a Proneurogenic Transcription Program. Neuron 84, 723–739. 10.1016/j.neuron.2014.10.022.

10. DeBoer, E.M., Azevedo, R., Vega, T.A., Brodkin, J., Akamatsu, W., Okano, H., Wagner, G.C., and Rasin, M.-R. (2014). Prenatal Deletion of the RNA-Binding Protein HuD Disrupts Postnatal Cortical Circuit Maturation and Behavior. J. Neurosci. 34, 3674–3686. 10.1523/JNEUROSCI.3703-13.2014.

11. Kraushar, M.L., Thompson, K., Wijeratne, H.R.S., Viljetic, B., Sakers, K., Marson, J.W., Kontoyiannis, D.L., Buyske, S., Hart, R.P., and Rasin, M.-R. (2014). Temporally defined neocortical translation and polysome assembly are determined by the RNA-binding protein Hu antigen R. Proc. Natl. Acad. Sci. U.S.A. 111. 10.1073/pnas.1408305111.

12. Popovitchenko, T., Park, Y., Page, N.F., Luo, X., Krsnik, Z., Liu, Y., Salamon, I., Stephenson, J.D., Kraushar, M.L., Volk, N.L., et al. (2020). Translational derepression of Elavl4 isoforms at their alternative 5′ UTRs determines neuronal development. Nat Commun 11, 1674. 10.1038/s41467-020-15412-8.

13. Kraushar, M.L., Viljetic, B., Wijeratne, H.R.S., Thompson, K., Jiao, X., Pike, J.W., Medvedeva, V., Groszer, M., Kiledjian, M., Hart, R.P., et al. (2015). Thalamic WNT3 Secretion Spatiotemporally Regulates the Neocortical Ribosome Signature and mRNA Translation to Specify Neocortical Cell Subtypes. Journal of Neuroscience 35, 10911–10926. 10.1523/JNEUROSCI.0601-15.2015.

14. Borisova, E., Newman, A.G., Couce Iglesias, M., Dannenberg, R., Schaub, T., Qin, B., Rusanova, A., Brockmann, M., Koch, J., Daniels, M., et al. (2024). Protein translation rate determines neocortical neuron fate. Nat Commun 15, 4879. 10.1038/s41467-024-49198-w.

15. Poulopoulos, A., Murphy, A.J., Ozkan, A., Davis, P., Hatch, J., Kirchner, R., and Macklis, J.D. (2019). Subcellular transcriptomes and proteomes of developing axon projections in the cerebral cortex. Nature 565, 356–360. 10.1038/s41586-018-0847-y.

16. Jung, H., Yoon, B.C., and Holt, C.E. (2012). Axonal mRNA localization and local protein synthesis in nervous system assembly, maintenance and repair. Nat Rev Neurosci 13, 308–324. 10.1038/nrn3210.

17. Campbell, D.S., and Holt, C.E. (2001). Chemotropic Responses of Retinal Growth Cones Mediated by Rapid Local Protein Synthesis and Degradation. Neuron 32, 1013–1026. 10.1016/S0896-6273(01)00551-7.

18. Davis, L., Dou, P., DeWit, M., and Kater, S. (1992). Protein synthesis within neuronal growth cones. J. Neurosci. 12, 4867–4877. 10.1523/JNEUROSCI.12-12-04867.1992.

19. Ming, G., Wong, S.T., Henley, J., Yuan, X., Song, H., Spitzer, N.C., and Poo, M. (2002). Adaptation in the chemotactic guidance of nerve growth cones. Nature 417, 411–418. 10.1038/nature745.

20. Shigeoka, T., Koppers, M., Wong, H.H.-W., Lin, J.Q., Cagnetta, R., Dwivedy, A., De Freitas Nascimento, J., Van Tartwijk, F.W., Ströhl, F., Cioni, J.-M., et al. (2019). On-Site Ribosome Remodeling by Locally Synthesized Ribosomal Proteins in Axons. Cell Reports 29, 3605–3619.e10. 10.1016/j.celrep.2019.11.025.

21. Froberg, J.E., Durak, O., and Macklis, J.D. (2023). Development of nanoRibo-seq enables study of regulated translation by cortical neuron subtypes, showing uORF translation in synaptic-axonal genes. Cell Reports 42, 112995. 10.1016/j.celrep.2023.112995.

22. Li, D., and Wang, J. (2020). Ribosome heterogeneity in stem cells and development. Journal of Cell Biology 219, e202001108. 10.1083/jcb.202001108.

23. Shi, Z., Fujii, K., Kovary, K.M., Genuth, N.R., Röst, H.L., Teruel, M.N., and Barna, M. (2017). Heterogeneous Ribosomes Preferentially Translate Distinct Subpools of mRNAs Genome-wide. Molecular Cell 67, 71–83.e7. 10.1016/j.molcel.2017.05.021.

24. Xue, S., Tian, S., Fujii, K., Kladwang, W., Das, R., and Barna, M. (2015). RNA regulons in Hox 5′ UTRs confer ribosome specificity to gene regulation. Nature 517, 33–38. 10.1038/nature14010.

25. Simsek, D., Tiu, G.C., Flynn, R.A., Byeon, G.W., Leppek, K., Xu, A.F., Chang, H.Y., and Barna, M. (2017). The Mammalian Ribo-interactome Reveals Ribosome Functional Diversity and Heterogeneity. Cell 169, 1051–1065.e18. 10.1016/j.cell.2017.05.022.

26. Deuerling, E., Gamerdinger, M., and Kreft, S.G. (2019). Chaperone Interactions at the Ribosome. Cold Spring Harb Perspect Biol 11, a033977. 10.1101/cshperspect.a033977.

27. Darnell, J.C., Van Driesche, S.J., Zhang, C., Hung, K.Y.S., Mele, A., Fraser, C.E., Stone, E.F., Chen, C., Fak, J.J., Chi, S.W., et al. (2011). FMRP Stalls Ribosomal Translocation on mRNAs Linked to Synaptic Function and Autism. Cell 146, 247–261. 10.1016/j.cell.2011.06.013.

28. Koppers, M., Özkan, N., Nguyen, H.H., Jurriens, D., McCaughey, J., Nguyen, D.T.M., Li, C.H., Stucchi, R., Altelaar, M., MacGillavry, H.D., et al. (2024). Axonal endoplasmic reticulum tubules control local translation via P180/RRBP1-mediated ribosome interactions. Developmental Cell 59, 2053–2068.e9. 10.1016/j.devcel.2024.05.005.

29. Tcherkezian, J., Brittis, P.A., Thomas, F., Roux, P.P., and Flanagan, J.G. (2010). Transmembrane Receptor DCC Associates with Protein Synthesis Machinery and Regulates Translation. Cell 141, 632–644. 10.1016/j.cell.2010.04.008.

30. Lai, T., Jabaudon, D., Molyneaux, B.J., Azim, E., Arlotta, P., Menezes, J.R.L., and Macklis, J.D. (2008). SOX5 controls the sequential generation of distinct corticofugal neuron subtypes. Neuron 57, 232–247. 10.1016/j.neuron.2007.12.023.

31. Molyneaux, B.J., Arlotta, P., Fame, R.M., MacDonald, J.L., MacQuarrie, K.L., and Macklis, J.D. (2009). Novel subtype-specific genes identify distinct subpopulations of callosal projection neurons. J Neurosci 29, 12343–12354. 10.1523/JNEUROSCI.6108-08.2009.

32. Galazo, M.J., Emsley, J.G., and Macklis, J.D. (2016). Corticothalamic Projection Neuron Development beyond Subtype Specification: Fog2 and Intersectional Controls Regulate Intraclass Neuronal Diversity. Neuron 91, 90–106. 10.1016/j.neuron.2016.05.024.

33. Sahni, V., Shnider, S.J., Jabaudon, D., Song, J.H.T., Itoh, Y., Greig, L.C., and Macklis, J.D. (2021). Corticospinal neuron subpopulation-specific developmental genes prospectively indicate mature segmentally specific axon projection targeting. Cell Rep 37, 109843. 10.1016/j.celrep.2021.109843.

34. Sahni, V., Itoh, Y., Shnider, S.J., and Macklis, J.D. (2021). Crim1 and Kelch-like 14 exert complementary dual-directional developmental control over segmentally specific corticospinal axon projection targeting. Cell Rep 37, 109842. 10.1016/j.celrep.2021.109842.

35. McKenna, W.L., Ortiz-Londono, C.F., Mathew, T.K., Hoang, K., Katzman, S., and Chen, B. (2015). Mutual regulation between Satb2 and Fezf2 promotes subcerebral projection neuron identity in the developing cerebral cortex. Proc. Natl. Acad. Sci. U.S.A. 112, 11702–11707. 10.1073/pnas.1504144112.

36. De León Reyes, N.S., Mederos, S., Varela, I., Weiss, L.A., Perea, G., Galazo, M.J., and Nieto, M. (2019). Transient callosal projections of L4 neurons are eliminated for the acquisition of local connectivity. Nat Commun 10, 4549. 10.1038/s41467-019-12495-w.

37. Lerner, E.A., Lerner, M.R., Janeway, C.A., and Steitz, J.A. (1981). Monoclonal antibodies to nucleic acid-containing cellular constituents: probes for molecular biology and autoimmune disease. Proc. Natl. Acad. Sci. U.S.A. 78, 2737–2741. 10.1073/pnas.78.5.2737.

38. Garden, G.A., Hartlage-Rübsamen, M., Rubel, E.W., and Bothwell, M.A. (1995). Protein Masking of a Ribosomal RNA Epitope Is an Early Event in Afferent Deprivation-Induced Neuronal Death. Molecular and Cellular Neuroscience 6, 293–310. 10.1006/mcne.1995.1023.

39. Younts, T.J., Monday, H.R., Dudok, B., Klein, M.E., Jordan, B.A., Katona, I., and Castillo, P.E. (2016). Presynaptic Protein Synthesis Is Required for Long-Term Plasticity of GABA Release. Neuron 92, 479–492. 10.1016/j.neuron.2016.09.040.

40. Schaeffer, J., Vilallongue, N., Decourt, C., Blot, B., El Bakdouri, N., Plissonnier, E., Excoffier, B., Paccard, A., Diaz, J.-J., Humbert, S., et al. (2023). Customization of the translational complex regulates mRNA-specific translation to control CNS regeneration. Neuron 111, 2881–2898.e12. 10.1016/j.neuron.2023.06.005.

41. Salehi, S., Zare, A., Prezza, G., Bader, J., Schneider, C., Fischer, U., Meissner, F., Mann, M., Briese, M., and Sendtner, M. (2023). Cytosolic Ptbp2 modulates axon growth in motoneurons through axonal localization and translation of Hnrnpr. Nat Commun 14, 4158. 10.1038/s41467-023-39787-6.

42. Zare, A., Salehi, S., Bader, J., Schneider, C., Fischer, U., Veh, A., Arampatzi, P., Mann, M., Briese, M., and Sendtner, M. (2024). hnRNP R promotes O-GlcNAcylation of eIF4G and facilitates axonal protein synthesis. Nat Commun 15, 7430. 10.1038/s41467-024-51678-y.

43. Sanz, E., Yang, L., Su, T., Morris, D.R., McKnight, G.S., and Amieux, P.S. (2009). Cell-type-specific isolation of ribosome-associated mRNA from complex tissues. Proc. Natl. Acad. Sci. U.S.A. 106, 13939–13944. 10.1073/pnas.0907143106.

44. Zhang, Y., O’Leary, M.N., Peri, S., Wang, M., Zha, J., Melov, S., Kappes, D.J., Feng, Q., Rhodes, J., Amieux, P.S., et al. (2017). Ribosomal Proteins Rpl22 and Rpl22l1 Control Morphogenesis by Regulating Pre-mRNA Splicing. Cell Rep 18, 545–556. 10.1016/j.celrep.2016.12.034.

45. Matzinger, M., Mayer, R.L., and Mechtler, K. (2023). Label-free sin-gle cell proteomics utilizing ultrafast LC and MS instrumentation: A valuable complementary technique to multiplexing. Proteomics 23, 2200162. 10.1002/pmic.202200162.

46. Bartsch, D., Kalamkar, K., Ahuja, G., Lackmann, J.-W., Hescheler, J., Weber, T., Bazzi, H., Clamer, M., Mendjan, S., Papantonis, A., et al. (2023). mRNA translational specialization by RBPMS presets the competence for cardiac commitment in hESCs. Sci. Adv. 9, eade1792. 10.1126/sciadv.ade1792.

47. Susanto, T.T., Hung, V., Levine, A.G., Chen, Y., Kerr, C.H., Yoo, Y., Oses-Prieto, J.A., Fromm, L., Zhang, Z., Lantz, T.C., et al. (2024). RAPIDASH: Tag-free enrichment of ribosome-associated proteins reveals composition dynamics in embryonic tissue, cancer cells, and macrophages. Molecular Cell 84, 3545–3563.e25. 10.1016/j.molcel.2024.08.023.

48. de la Cruz, J., Karbstein, K., and Woolford, J.L. (2015). Functions of ribosomal proteins in assembly of eukaryotic ribosomes in vivo. Annu Rev Biochem 84, 93–129. 10.1146/annurev-biochem-060614-033917.

49. Van Den Heuvel, J., Ashiono, C., Gillet, L.C., Dörner, K., Wyler, E., Zemp, I., and Kutay, U. (2021). Processing of the ribosomal ubiqui-tin-like fusion protein FUBI-eS30/FAU is required for 40S maturation and depends on USP36. eLife 10, e70560. 10.7554/eLife.70560.

50. Oughtred, R., Rust, J., Chang, C., Breitkreutz, B., Stark, C., Willems, A., Boucher, L., Leung, G., Kolas, N., Zhang, F., et al. (2021). The BI-OGRID database: A comprehensive biomedical resource of curated protein, genetic, and chemical interactions. Protein Science 30, 187–200. 10.1002/pro.3978.

51. Cho, N.H., Cheveralls, K.C., Brunner, A.-D., Kim, K., Michaelis, A.C., Raghavan, P., Kobayashi, H., Savy, L., Li, J.Y., Canaj, H., et al. (2022). OpenCell: Endogenous tagging for the cartography of human cellular organization. Science 375, eabi6983. 10.1126/science.abi6983.

52. Gemmer, M., Chaillet, M.L., Van Loenhout, J., Cuevas Arenas, R., Vismpas, D., Gröllers-Mulderij, M., Koh, F.A., Albanese, P., Scheltema, R.A., Howes, S.C., et al. (2023). Visualization of translation and protein biogenesis at the ER membrane. Nature 614, 160–167. 10.1038/s41586-022-05638-5.

53. Schaffer, T.B., Smith, J.E., Cook, E.K., Phan, T., and Margolis, S.S. (2018). PKCε Inhibits Neuronal Dendritic Spine Development through Dual Phosphorylation of Ephexin5. Cell Reports 25, 2470–2483.e8. 10.1016/j.celrep.2018.11.005.

54. Sen, A., Hongpaisan, J., Wang, D., Nelson, T.J., and Alkon, D.L. (2016). Protein Kinase Cϵ (PKCϵ) Promotes Synaptogenesis through Membrane Accumulation of the Postsynaptic Density Protein PSD-95. Journal of Biological Chemistry 291, 16462–16476. 10.1074/jbc.M116.730440.

55. Genuth, N.R., and Barna, M. (2018). The Discovery of Ribosome Heterogeneity and Its Implications for Gene Regulation and Organismal Life. Molecular Cell 71, 364–374. 10.1016/j.molcel.2018.07.018.

56. Fusco, C.M., Desch, K., Dörrbaum, A.R., Wang, M., Staab, A., Chan, I.C.W., Vail, E., Villeri, V., Langer, J.D., and Schuman, E.M. (2021). Neuronal ribosomes exhibit dynamic and context-dependent exchange of ribosomal proteins. Nat Commun 12, 6127. 10.1038/s41467-021-26365-x.

57. Ladd, A.N., Charlet-B., N., and Cooper, T.A. (2001). The CELF Family of RNA Binding Proteins Is Implicated in Cell-Specific and Developmentally Regulated Alternative Splicing. Mol Cell Biol 21, 1285–1296. 10.1128/MCB.21.4.1285-1296.2001.

58. Ladd, A.N., Nguyen, N.H., Malhotra, K., and Cooper, T.A. (2004). CELF6, a Member of the CELF Family of RNA-binding Proteins, Regulates Muscle-specific Splicing Enhancer-dependent Alternative Splicing. Journal of Biological Chemistry 279, 17756–17764. 10.1074/jbc.M310687200.

59. Horb, L.D., and Horb, M.E. (2010). BrunoL1 regulates endoderm proliferation through translational enhancement of cyclin A2 mRNA. Developmental Biology 345, 156–169. 10.1016/j.yd-bio.2010.07.005.

60. Pérez-González, A., Pazo, A., Navajas, R., Ciordia, S., Rodriguez-Frandsen, A., and Nieto, A. (2014). hCLE/C14orf166 Associates with DDX1-HSPC117-FAM98B in a Novel Transcription-Dependent Shuttling RNA-Transporting Complex. PLoS ONE 9, e90957. 10.1371/journal.pone.0090957.

61. Kanai, Y., Dohmae, N., and Hirokawa, N. (2004). Kinesin Transports RNA. Neuron 43, 513–525. 10.1016/j.neuron.2004.07.022.

62. Edmondson, R.D., Vondriska, T.M., Biederman, K.J., Zhang, J., Jones, R.C., Zheng, Y., Allen, D.L., Xiu, J.X., Cardwell, E.M., Pisano, M.R., et al. (2002). Protein Kinase C ε Signaling Complexes Include Metabolism- and Transcription/Translation-related Proteins. Molecular & Cellular Proteomics 1, 421–433. 10.1074/mcp.M100036-MCP200.

63. Gorski, J.A., Talley, T., Qiu, M., Puelles, L., Rubenstein, J.L.R., and Jones, K.R. (2002). Cortical Excitatory Neurons and Glia, But Not GABAergic Neurons, Are Produced in the Emx1-Expressing Line-age. J. Neurosci. 22, 6309–6314. 10.1523/JNEURO-SCI.22-15-06309.2002.

64. Madisen, L., Zwingman, T.A., Sunkin, S.M., Oh, S.W., Zariwala, H.A., Gu, H., Ng, L.L., Palmiter, R.D., Hawrylycz, M.J., Jones, A.R., et al. (2010). A robust and high-throughput Cre reporting and characterization system for the whole mouse brain. Nat Neurosci 13, 133–140. 10.1038/nn.2467.

65. Catapano, L.A., Arnold, M.W., Perez, F.A., and Macklis, J.D. (2001). Specific Neurotrophic Factors Support the Survival of Cortical Projection Neurons at Distinct Stages of Development. J. Neurosci. 21, 8863–8872. 10.1523/JNEUROSCI.21-22-08863.2001.

66. Engmann, A.K., Hatch, J.J., Nanda, P., Veeraraghavan, P., Ozkan, A., Poulopoulos, A., Murphy, A.J., and Macklis, J.D. (2022). Neuronal subtype-specific growth cone and soma purification from mammalian CNS via fractionation and fluorescent sorting for subcellular analyses and spatial mapping of local transcriptomes and proteomes. Nat Protoc 17, 222–251. 10.1038/s41596-021-00638-7.

67. Käll, L., Storey, J.D., and Noble, W.S. (2008). Non-parametric estimation of posterior error probabilities associated with peptides identified by tandem mass spectrometry. Bioinformatics 24, i42–i48. 10.1093/bioinformatics/btn294.

68. Thomas, P.D., Ebert, D., Muruganujan, A., Mushayahama, T., Albou, L., and Mi, H. (2022). PANTHER : Making genome-scale phylogenetics accessible to all. Protein Science 31, 8–22. 10.1002/pro.4218.

69. Mi, H., Muruganujan, A., Huang, X., Ebert, D., Mills, C., Guo, X., and Thomas, P.D. (2019). Protocol Update for large-scale genome and gene function analysis with the PANTHER classification system (v.14.0). Nat Protoc 14, 703–721. 10.1038/s41596-019-0128-8.

70. Zhang, X., Smits, A.H., Van Tilburg, G.B., Ovaa, H., Huber, W., and Vermeulen, M. (2018). Proteome-wide identification of ubiquitin interactions using UbIA-MS. Nat Protoc 13, 530–550. 10.1038/nprot.2017.147.

71. Schindelin, J., Arganda-Carreras, I., Frise, E., Kaynig, V., Longair, M., Pietzsch, T., Preibisch, S., Rueden, C., Saalfeld, S., Schmid, B., et al. (2012). Fiji: an open-source platform for biological-image analysis. Nat Methods 9, 676–682. 10.1038/nmeth.2019.

